# Bidirectional epigenetic editing reveals hierarchies in gene regulation

**DOI:** 10.1101/2022.11.15.516658

**Authors:** Naomi M. Pacalin, Quanming Shi, Kevin R. Parker, Howard Y. Chang

## Abstract

CRISPR perturbations are valuable tools for studying the functional effects of the genome. However, existing methods are limited in their utility for studying noncoding elements and genetic interactions. Here, we develop a system for bidirectional epigenetic editing (CRISPRai), in which orthogonal activating (CRISPRa) and repressive (CRISPRi) perturbations are applied simultaneously to multiple loci the same cell. We developed dual-gRNA-capture single-cell Perturb-seq to study the established interaction between SPI1 and GATA1, two hemopoietic lineage transcription factors, and discovered novel context-specific regulation modes for co-regulated genes. Extending CRISPRai to noncoding elements, we addressed how multiple enhancers interact to modulate expression of a shared target gene, Interleukin-2, in T cells. We found that enhancer function was primarily additive and enabled fine-tuning of gene expression, yet a clear hierarchy existed among enhancers in strength of gene expression control. The promoter was dominant over most enhancers in controlling gene expression; however, a small subset of enhancers exhibited strong functional effects, or gatekeeper function, and could turn off the gene despite promoter activation. Integration of these functional data with histone ChIP-seq and TF motif enrichment suggests the existence of multiple modes of enhancer-mediated gene regulation. Our method, CRISPRai for bidirectional epigenetic editing, provides an approach for identifying novel genetic interactions that may be overlooked when studied without bidirectional perturbations and can be applied to both genes and noncoding elements.

## INTRODUCTION

Programmable epigenetic editing, specifically CRISPRa^1-5^ and CRISPRi^6,7^, are valuable tools for uncovering functional effects of genes and noncoding elements such as enhancers^8-15^. Dual CRISPR perturbations, in which two genes are perturbed simultaneously, are uniquely able to identify genetic interactions and epistasis, which in turn enables the rapid mapping of genetic pathways^16-20^. Previously, large-scale dual CRISPR perturbation screens employed CRISPRa and CRISPRko^18,21-23^, which introduced dsDNA breaks via Cas9 nuclease cutting, limiting multiplexing capacity due to risk of translocations and cellular toxicity from DNA damage and indels^24-29^ and limiting applicability to noncoding elements for which small indels typically do not alter function. More recently, methods for bidirectional perturbations of two loci simultaneously, including paired CRISPRa and CRISPRi, have been developed, but have only been applied to non-mammalian cells, are transient, or targeted to a few genes^30-35^. These studies have not demonstrated the feasibility of bidirectional systems for understanding human genetic interactions on a transcriptome-wide scale, for pooled screens with a massively parallel single-cell readout, or for the investigation of noncoding elements. Multiplexed CRISPRi can address noncoding element epistasis^36^, but may be limited to elements that are contemporaneously active in the cell type being studied. Epigenetic perturbations are key for studying functional effects of noncoding elements such as enhancers in their endogenous locus because enhancer functionality is likely mediated through structural chromatin contacts, histone modifications, transcription factor (TF) requirement, and other effects^37-44^. Additionally, comprehensive investigation of interactions between genes requires versatile bidirectional perturbation tools in order to study the complete range of context-specific genetic interactions^8,45-47^.

To broaden the toolkit for studying genes and noncoding elements and to enable investigation of context-specific genetic interactions, we developed CRISPRai, a system for simultaneous bidirectional epigenetic editing of two loci in a single cell. We use orthogonal activating (CRISPRa) and repressive (CRISPRi) perturbations to perturb two distinct genomic loci simultaneously, effectively turning one element “on” and the other “off”, in order to study how two genetic elements functionally interact. We developed dual-gRNA-capture Perturb-seq, a method for single-cell transcriptome profiling coupled with CRISPR guide RNA (gRNA) readout^48-52^, to demonstrate bidirectional perturbation of CRISPR target genes. Investigation of the SPI1 and GATA1 genetic interaction^53-56^ through single-cell transcriptomics confirmed regulatory effects of these TFs on previously known downstream targets and determined novel characteristics of their interaction on co-regulated genes. In addition, we applied CRISPRai to investigate how multiple enhancers for a given gene interact to control target gene expression. We used the Interleukin-2 (*IL2*) locus and *IL2* gene expression in activated Jurkat T cells as a model system. Our results demonstrated that the promoter generally supersedes enhancers in the hierarchy of target gene expression control, that bidirectional pairwise epigenetic enhancer editing enables tuning of target gene expression, and that most enhancers combine to act in an additive manner. Further, we identified the presence of gatekeeper enhancers which strongly influenced target gene expression, were dominant over other enhancers, and could act as an off-switch for target gene expression. Our method, CRISPRai, highlights novel genetic interactions for both genes and noncoding elements, particularly for genetic elements whose function may be overlooked in a one-directional perturbation context, and broadens the toolkit for investigating the functional effects of the genome.

## RESULTS

### CRISPRai system for bidirectional epigenetic editing in individual cells

We developed a system for bidirectional epigenetic editing that enables activation and repression of two distinct loci simultaneously in a single cell, which can be applied to study genes and enhancers (**Fig. 1a, Supplementary Fig. S1a-i**). Our system comprises Tet-On doxycycline-inducible (dox) CRISPRa and CRISPRi machinery (CRISPRai) and leverages two orthogonal species of catalytically dead Cas9 (dCas9) as programmable DNA binding platforms. We express activator-fused dCas9 from*S. aureus* (VPR-dSaCas9) and repressor-fused dCas9 from *S. pyogenes* (dSpCas9-KRAB) to achieve species-specific recognition through unique gRNA scaffold sequences^57,58^, thus achieving two distinct perturbations at two different loci in the same cell (**Fig. 1a-b, Supplementary Fig. S1a-i**). After generating stable K562 (**Supplementary Fig. S1a-e**) and Jurkat (**Supplementary Fig. S1f-i**) CRISPRai cell lines, we validated the system using bulk assays. We confirmed construct expression, inducibility, CRISPR perturbation dependence on dCas9 expression level (**Supplementary Fig. S1a,b**), demonstrated single and double bidirectional perturbations for several genes ranging from −2.5 to 13 log2FC gene expression change (**Supplementary Fig. S1c,f-h**), and ensured stable construct expression over a 14-20 day timeline of dCas9 induction (**Supplementary Fig. S1d,e,i**). The magnitude of gene expression perturbation was comparable in the single and double perturbations (**Supplementary Fig. S1c**).

**Fig. 1.**
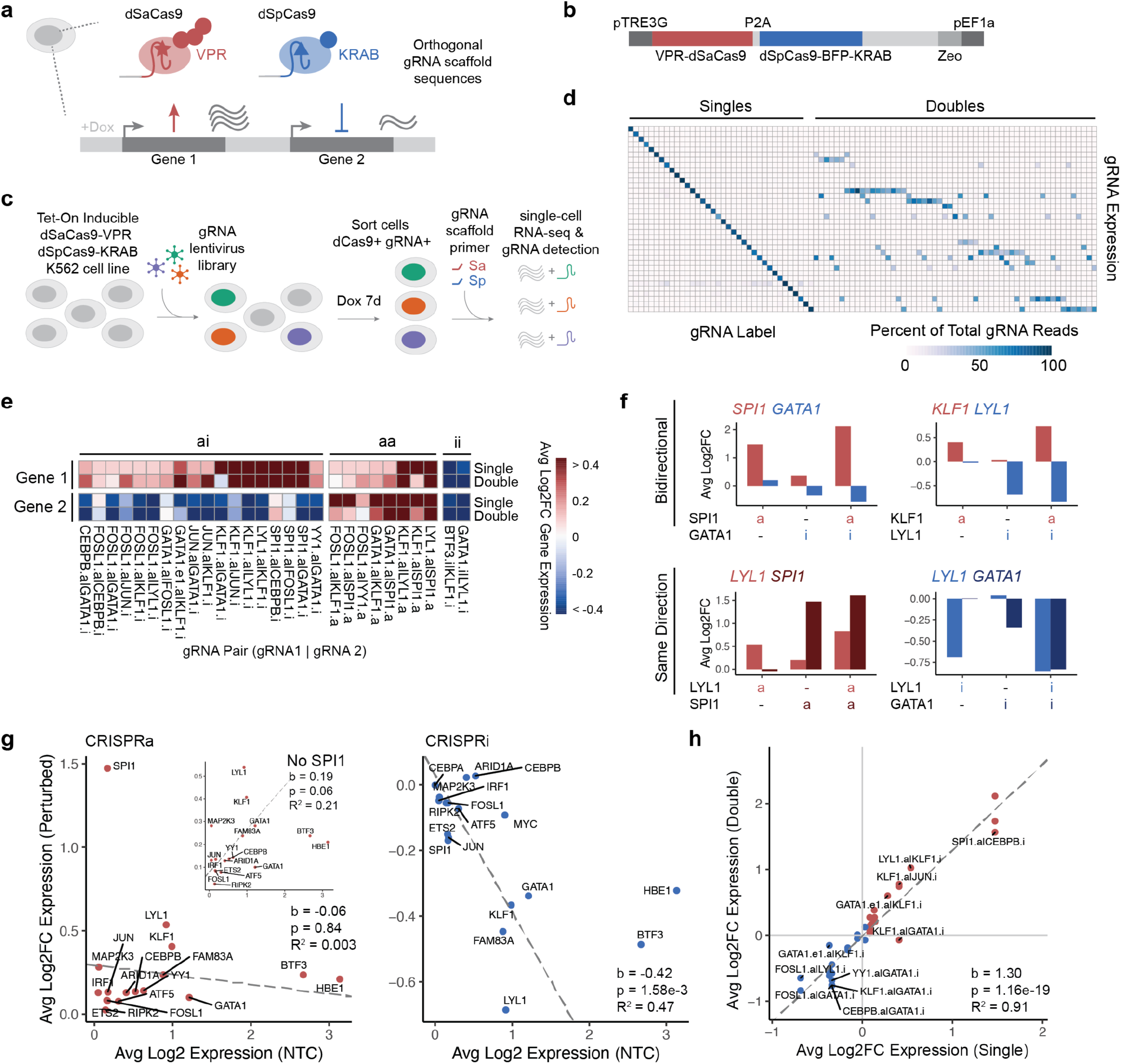
CRISPRai system for bidirectional epigenetic editing in individual cells. Single-cell CRISPRai screen in K562 cells. (a) Schematic of CRISPRai system. (b) Schematic of CRISPRai construct. (c) Screen workflow for scRNA-seq and gRNA detection. (d) Heatmap showing gRNA detection (rows) by groups of cells labeled for that gRNA (columns) in single and double perturbations (singles and doubles, respectively). Shown as percent of total gRNA reads. Cells are grouped and averaged by gRNA call. (e) Heatmap summary of CRISPRai perturbations for a panel of gene pairs. Data shows average log2FC gene expression for each pair of target genes (columns) in cells receiving either a single or double perturbation (rows). Column annotations show type of perturbation applied (ai, aa, ii). (f) Examples of average log2FC gene expression in single and double perturbations for indicated target gene pairs with ai, aa, or ii perturbations. (g) Perturbation strength (average log2FC gene expression in perturbed cells) as a function of baseline target gene expression level in NTC cells (average log2 gene expression in NTC cells), for (left) CRISPRa and (right) CRISPRi. (h) Correlation of perturbation strength (average log2FC gene expression) in single and double perturbations for a given target gene, labeled with perturbation received. e-h differential expression tests performed relative to cells with NTC gRNAs.

We next developed dual-gRNA-capture Perturb-seq to study gene-gene interactions with a single-cell transcriptome readout in K562 cells. We introduced a lentiviral pool of 82 single, 22 double, and 12 non-targeting control (NTC) gRNAs containing selected combinations of single and double perturbations against a panel of 19 lineage-relevant TFs, chromatin remodelers, and proto-oncogenes, with 2 gRNAs per gene (**Supplementary Fig. S2a, Supplementary Table S1**,**S2**). We then detected the transcriptome and gRNA expression in single cells (**Fig. 1c**). For gRNA detection, we extended previous methods of droplet-based direct gRNA sequence detection^59,60^ to enable dual gRNA detection for both dCas9 species gRNA scaffolds by spiking in oligos complementary to the gRNA scaffold region in the reverse transcription reaction using 5’ RNA capture; two oligos were added, one oligo for each dCas9 species gRNA scaffold. Single and double gRNAs were accurately detected, allowing single cells to be grouped based on the identity of their perturbations (**Fig. 1d**). We captured a total of 24,661 cells, 14,086 cells with single perturbations, 6,631 cells with double perturbations, and 3,944 cells with NTCs (**Supplementary Fig. S3a**). We first investigated whether the gRNA target genes were perturbed in the desired direction, if bidirectional control of two genes was possible in the single-cell context, and if this control was possible for a breadth of genes. We found that the system enabled consistent bidirectional expression changes in both target genes in all double combinations, with the log2FC gene expression increasing or decreasing as expected in each condition, ranging from −0.85 to +2.11 log2FC gene expression change (**Fig. 1e,f, Supplementary Fig. S3b**). In addition to CRISPRai combinations, we demonstrated the expected behavior for CRISPRaa and CRISPRii combinations as well; the CRISPRai system allows for CRISPRaa and CRISPRii combinations by delivering two gRNAs with the same gRNA scaffold to a given cell, which consequently directs the gRNAs to the same dCas9 species and leads to dual same-direction perturbations. The perturbed expression changes were statistically significant in both the single and double perturbations and spanned a range of log2FC across different genes (**Fig. 1g, Supplementary Fig. S3b**). We found that different genes had variable susceptibility to perturbation. For example, *SPI1*, which has low expression in K652s, was highly responsive to activation, but not repression (**Fig. 1g, Supplementary Fig. S3b**). *GATA1*, a highly expressed gene in K562s, was highly responsive to repression, and less so to activation. *LYL1* and *KLF1*, both with mid expression in K562s, were both responsive to activation and repression (**Fig. 1g, Supplementary Fig. S3b)**.

We then investigated the characteristics of bidirectional editing across the genes in the panel to understand its utility in perturbing other genes in a broader context. In general, the double and single log2FC target gene expression in perturbed versus non-perturbed cells was highly correlated (R^2^=0.91, b=1.30, p=<1.16e-19, **Fig. 1h**). This further supports the orthogonality of the system and indicates that the double perturbation condition does not dilute the perturbative capacity of the individual perturbations in the pair, an issue that can arise with multiplexed CRISPR perturbations^61-63^. Further, baseline gene expression inversely correlated with perturbation strength for CRISPRi; highly expressed genes could be repressed with high log2FC gene expression, whereas lowly expressed genes can only be slightly repressed (R^2^=0.47, b=-0.42, p=1.58e-3, **Fig. 1g right**). This is expected because setting gene expression to zero acts a lower limit on gene repression potential; it is not possible to further repress a gene that is already silent. In contrast, baseline gene expression and strength of CRISPRa did not have a clear relationship; this may be due to gene-specific regulatory mechanisms, different chromatin modifying functions of VPR and KRAB, or that there is a less stringent upper limit on gene activation (R^2^=0.003, b=-0.06, p=0.84, **Fig. 1g left**). Additionally, double perturbations tended to have more differentially expressed (DE) genes than single perturbations (**Supplementary Fig. S3c**). Overall, bidirectional editing with CRISPRai allows us to examine the full set of potential regulators irrespective of their current gene expression state in the cell type being studied.

### CRISPRai reveals context-specific genetic interactions

Pairwise CRISPR perturbations can identify genetic interactions between genes involved in similar biological pathways^16-20,23^, and CRISPR screens with high-content readouts enable investigation of the global regulatory effects of a given gene, including identification of downstream target genes and regulatory gene modules controlled by the perturbed gene^48-52,60^. We applied bidirectional epigenetic editing to investigate genetic interactions. We reasoned that since certain genes have greater responsiveness to either activation or repression, in some cases bidirectional control (CRISPRai) is required to study a given pair of interacting genes and that consequently bidirectional control can reveal novel functionality that would otherwise be missed using a toolkit that only enables pairwise unidirectional perturbations (i.e. CRISPRaa or CRISPRii). A good example of this phenomenon is *SPI1* and *GATA1*, as each gene is strongly responsive to activation or repression, respectively (**Fig. 1e-f**). SPI1 and GATA1 are pivotal hematopoietic-specific TFs that are essential for myeloid and erythroid lineage development, and they are known to interact and inhibit each other’s function^53-56^.

We first investigated the global transcriptome-wide effects of all combinations of SPI1 and GATA1 perturbations included in the screen, including single and double perturbation combinations, in 437 cells. Notably, using single-cell RNA-seq (scRNA-seq) data, cells clustered according to the detected gRNAs and clusters were ordered according to the perturbation; double perturbations (doubles) clustered between the corresponding single perturbations (singles), demonstrating a gradient in transcriptomic signature resulting from the perturbations (**Fig. 2a, Supplementary Fig. S4a-d**). Additionally, the transcriptomes in doubles were most highly correlated with the transcriptomes of the corresponding singles (**Fig. 2b, Supplementary Fig. S4d**). The set of DE genes (relative to NTC gRNAs) in the bidirectional SPI1-a|GATA1-i double (ai), was composed of two subsets that were shared by the corresponding singles, and a third double-specific subset of 70 genes (**Fig. 2c, Supplementary Fig. S5a-c**). The upregulated DE genes for each perturbation condition relative to NTC were enriched for relevant biological process GO terms, including myeloid cell activation, with the double perturbation being most significantly enriched (**Fig. 2d**). Further, the subset of upregulated DE genes that was unique to the double perturbation and not present in the single perturbations was similarly enriched for relevant biological process GO terms (**Supplementary Fig. S5c**).

**Fig. 2.**
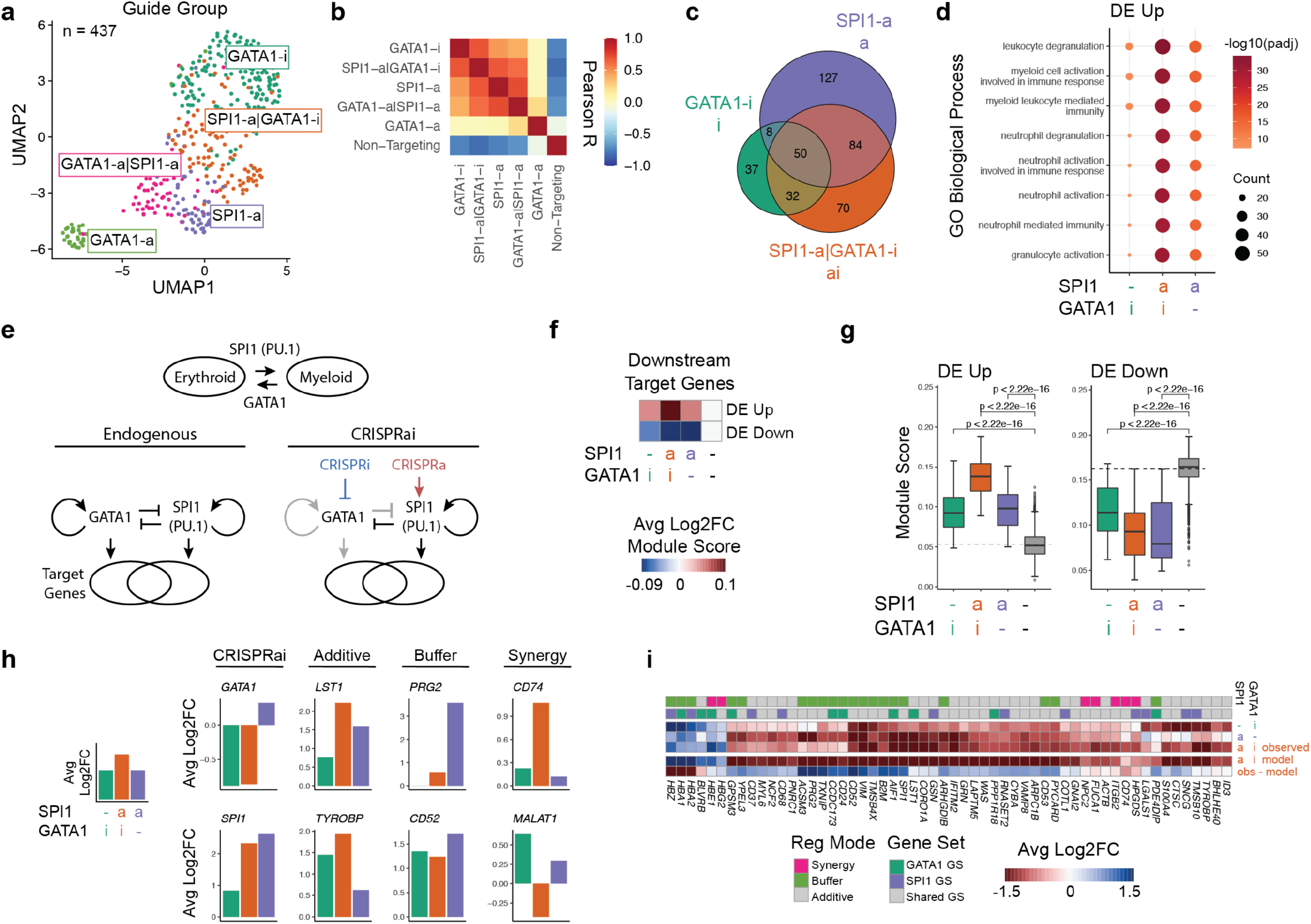
CRISPRai reveals context-specific genetic interaction for SPI1 and GATA1. Data from single-cell CRISPRai K562 screen in **Fig. 1**. UMAP representation of single-cell transcriptomes of perturbation-driven cells (**Supplementary Fig. S4c**) in the SPI1-GATA1 perturbation set. (b) Pearson correlation of single-cell transcriptomes over all genes by gRNA group. (c) Overlap of DE genes across gRNA groups. (d) Biological process GO term enrichment by gRNA group for DE genes upregulated in perturbed cells relative to NTC, binned by perturbation. Gene count >20 in at least one group, top 8 most significant terms shown. (e) Schematic of known SPI1-GATA1 genetic interaction and predicted changes under CRISPRai perturbation. (f) Average log2FC module score for the combined ENCODE TF target gene set^64–66^ for SPI1 and GATA1, partitioned by up or down regulation in the double perturbation. Module score is a measure of average expression of a gene set. (g) Same data as f. (h) Average log2FC gene expression of *SPI1* and *GATA1* and several of their ENCODE annotated downstream target genes, showing examples of each regulatory mode. (i) Heatmap of average log2FC gene expression by gRNA group of top DE genes from ENCODE TF target gene sets for SPI1 and GATA1 (Adj_p_value < 0.07, Abs(logFC) > 1, minimum % cells expressing gene > 10%). Expected double log2FC is calculated from additive model of singles, difference between observed and expected double (observed-expected) is shown. Gene set membership and regulatory mode of SPI1 and GATA1 on the gene is annotated, synergy diff > 0.1 (greater magnitude than expected or opposite sign than expected), buffer diff > 0.8 (lower magnitude than expected). c,d,h,i significance cutoffs for DE genes are abs(log2FC) > 0.5, p_adj < 0.05 unless noted otherwise, DE gene testing for each gRNA group is against NTC.

To confirm the utility of the bidirectional epigenetic editing system, we asked if known downstream target genes of SPI1 and GATA1 followed expected changes in each of the perturbation conditions aspredicted by the established SPI1-GATA1 functional relationship (**Fig. 2e**). Since SPI1 and GATA1 exhibit opposing and antagonistic effects on the myeloid and erythroid lineages, we hypothesized that known downstream target gene sets would have a more extreme gene expression change in the double (ai) relative to the single perturbations (a or i). We used the set of annotated target genes for these two TFs from ENCODE, which is determined by TF ChIP-seq^64-66^, and partitioned the gene sets based on up or down regulation in the double perturbation. As expected, the average expression of known target genes was more extreme in the double (ai) than the singles (a or i) (**Fig. 2f,g**). Further, this pattern persisted after partitioning the gene sets based on identity of TF regulator: GATA1 only, SPI1 only, or shared (**Supplementary Fig. S5d,e**). We validated this regulatory pattern on TF target gene sets from a different database (MSigDb)^67,68^ and saw similar results (**Supplementary Fig. S5f-h**). Additionally, we confirmed that the set of statistically significant DE genes in the double perturbation was highly overlapping with annotated SPI1 and GATA1 target genes sets (**Supplementary Fig. S5i,j**).

We then investigated if there were downstream target genes whose regulation was strongly influenced by the interaction between SPI1 and GATA1. We used an additive model of gene regulation that has previously been used for pairwise CRISPR perturbations^49,50,69^, in which the expected gene expression in the double perturbation is estimated from the sum of the single perturbations. Among genes that were differentially expressed in the perturbed cells and genes annotated as being regulated by SPI1, GATA1, or both^64-66^, we identified target genes that were governed by buffering, synergistic, or additive regulation (magnitude of log2FC less than, greater than, or equal to expected) (**Fig. 2h,i**). The regulatory mode for a given gene was independent of the annotated gene set membership, suggesting that the interaction between SPI1 and GATA1 is more complex than previously thought, that the set of target genes for each TF may be broader than previously annotated, that indirect regulatory effects can lead to dramatic gene expression changes, or that extent of co-regulation of each target gene by SPI1 and GATA1 is gene-specific (**Fig. 2i**).

Several genes under buffering regulation were key disease-associated genes. These include *CD52*, encoding a monoclonal antibody target for treatment of chronic lymphocytic leukemia^70^, *ACSM3*, a known tumor suppressor gene implicated in ovarian cancer^71^, and *PRG2*, a gene implicated in certain inflammatory gastrointestinal diseases^72,73^ (**Fig. 2i**). Several genes under synergistic regulation were also implicated in human diseases, including *CD74*, encoding a monoclonal antibody target for treatment of leukemia^74^, and *ITGB2*, a gene identified as prognostic in some cancers^75^ (**Fig. 2i**). Overall, this suggests that bidirectional epigenetic editing identifies regulatory modes for important, disease-associated genes.

### CRISPRai defines hierarchies in transcriptional regulation between promoters and enhancers

After demonstrating the efficacy of the CRISPRai system for investigating trans-regulatory effects and gene-gene interactions, we next extended our method to investigate cis-regulatory effects by studying enhancer-promoter (E-P, E-TSS) and enhancer-enhancer (E-E) interactions. Previous studies have shown that enhancer activity on target gene expression is governed by several factors, including distance to transcriptional start site (TSS) and enhancer strength, and that some enhancers may have redundant function or contribute highly to target gene expression levels^37,44,76,77^. However, it is unknown how multiple enhancers may interact to control target gene expression or how enhancers interact differentially with the TSS. We applied CRISPRai to study the regulatory landscape of a given gene and investigate how different enhancers interact with the promoter and with other enhancers in the locus.

We applied bidirectional epigenetic perturbations to 10 predicted enhancers and the promoter of the Interleukin-2 (*IL2*) gene in human Jurkat T cells (**Fig. 3a**). *IL2* is a key cytokine gene with a relatively large set of predicted enhancers, spanning a 2.4 Mb range^42^, making it an interesting gene for which to study enhancer interactions in both short and long range (**Fig. 3a, Supplementary Fig. S7f**). Enhancers with high predicted enhancer scores for the *IL2* gene in the Activity-by-Contact (ABC) Model^41,42^, a model that uses a combination of genomic datasets for gene-specific enhancer prediction, were selected for the screen. Some selected enhancers exhibited strong enhancer-related ChIP-seq signal, while others did not (**Fig. 3a**). Using our Jurkat T cell line expressing inducible CRISPRai machinery (**Supplementary Fig. S1f-i**), we introduced a lentiviral pool of 576 gRNA pairs (484 double, 88 single, 4 NTC gRNA pairs, **Fig. 3b, Supplementary Table S3**). The gRNA pool contained all CRISPRa and CRISPRi single perturbations (singles) and all CRISPRai pairwise combinations (doubles) for each enhancer and the TSS, as well as NTCs (**Supplementary Fig. S2b**). After 6 days of CRISPRai induction, the cells were activated to induce cytokine expression (**Supplementary Fig. S6a-c**), sorted for cytokine positive and negative populations (∼22% IL2+, ∼0.1% IFNG+, ∼1% IL2+IFNG+, **Fig. 3c left**), gRNA enrichment libraries were constructed (**Fig. 3b, Supplementary Fig. S7a-e**), and all CRISPRa and CRISPRi pairs were examined (**Fig. 3d, Supplementary Fig. S7g-i**). In addition to the *IL2* locus gRNA pool, we also performed an Interferon-gamma (*IFNG*) locus screen with 625 gRNA pairs (**Fig. 3c right, Supplementary Fig. S8a-i, Supplementary Table S4**), but due to low IFNG+ cell number, we only focused on TSS-E interactions for *IFNG* (**Fig. 3j,k**). First, we investigated how the TSS interacts with the enhancers globally. We compared log2FC gRNA enrichment in IL2+ versus cytokine-negative populations (IL2+ / NEG) and found that TSS-E interactions followed a largely additive relationship, whereby the expected double (log2FC gRNA enrichment z-scores in each corresponding single were summed) was highly correlated with the observed double (R^2^=0.91, b=0.97, p=<2e-16, **Fig. 3e**). The theoretical (x=y) and actual fits were highly overlapping (**Fig. 3e**). Log2FC gRNA enrichment z-scores ranged from ∼ −20 with CRISPRi to ∼ +7.5 with CRISPRa (**Fig. 3e**), and the true gRNA enrichment fold change (FC) ranged from 0.48 to 1.54 (not shown). We noted that some enhancers had strong functional effects while others had weaker functional effects in single perturbations (**Fig. 3d**). Further, this additive model held for all TSS-E doubles regardless of TSS perturbation strength in a subsequent validation gRNA pool (**Supplementary Fig. S7j**). We binned TSS-E doubles based on strength of TSS gRNA in singles, leveraging the natural variation in strength of TSS gRNAs. We then calculated residuals of the double from the global additive model fit, to obtain a measure of how different the expected and observed doubles were. The distribution of residuals of TSS-E doubles centered on zero and remained effectively constant across TSS perturbation strength (**Supplementary Fig. S7j**), demonstrating that the additive model holds regardless of perturbation strength. We then investigated how each individual enhancer interacts with the TSS (**Fig. 3d,f-k**). In general, the TSS exhibited clear hierarchy over enhancers (**Fig. 3g-k, Supplementary Fig. S7h, Supplementary Fig. S8i, Supplementary Fig. S9h**). In other words, the TSS perturbation was functionally dominant over enhancer perturbations and therefore acted as the driver of the gRNA enrichment level and consequently target gene expression. Silencing the promoter prevented most of the activated enhancers from activating *IL2* or *IFNG* (**Fig. 3h,k**). Conversely, activating the promoter largely bypassed the effect of enhancer silencing to activate gene expression (**Fig. 3g,j**). However, two enhancers were exceptions to this trend, namely E4 and E6 for *IL2* (**Fig. 3g,h**). Repression of these two enhancers individually was sufficient to counteract TSS activation and significantly reduce target gene expression (**Fig. 3g**). However, in the reverse condition, TSS inhibition with enhancer activation, ability to counteract TSS inhibition was minimal for E4 and not observed for E6 (**Fig. 3h**). This behavior suggests that E4 and E6 may act as “off-switches” or “gatekeepers” for *IL2* expression, in that that they are strong functional enhancers capable of overcoming the perturbation applied to the TSS. Examination of these gRNA pairings with true FC rather than log2FC z-scores revealed the same patterns (not shown). For *IFNG*, log2FC gRNA enrichment z-scores ranged from ∼ −3.5 for CRISPRi to ∼ +14 with CRISPRa, and TSS activation held hierarchy over enhancer inhibition for all enhancers, with E4 inhibition minimally counteracting TSS activation (**Fig. 3j**). In the reverse condition, TSS repression with enhancer activation, E7 strongly counteracted TSS repression, suggesting that E7 may act as an “on-switch” for *IFNG* when the TSS is repressed (**Fig. 3k**). We confirmed that no gRNAs had off-targets within 2kb of the corresponding gene’s TSS using Cas OFFinder^78^, indicating that the observed enhancer behavior is not due to off-target effects (parameters: mismatches <=4, DNA and RNA bulge sizes <=1) (not shown).

**Fig. 3.**
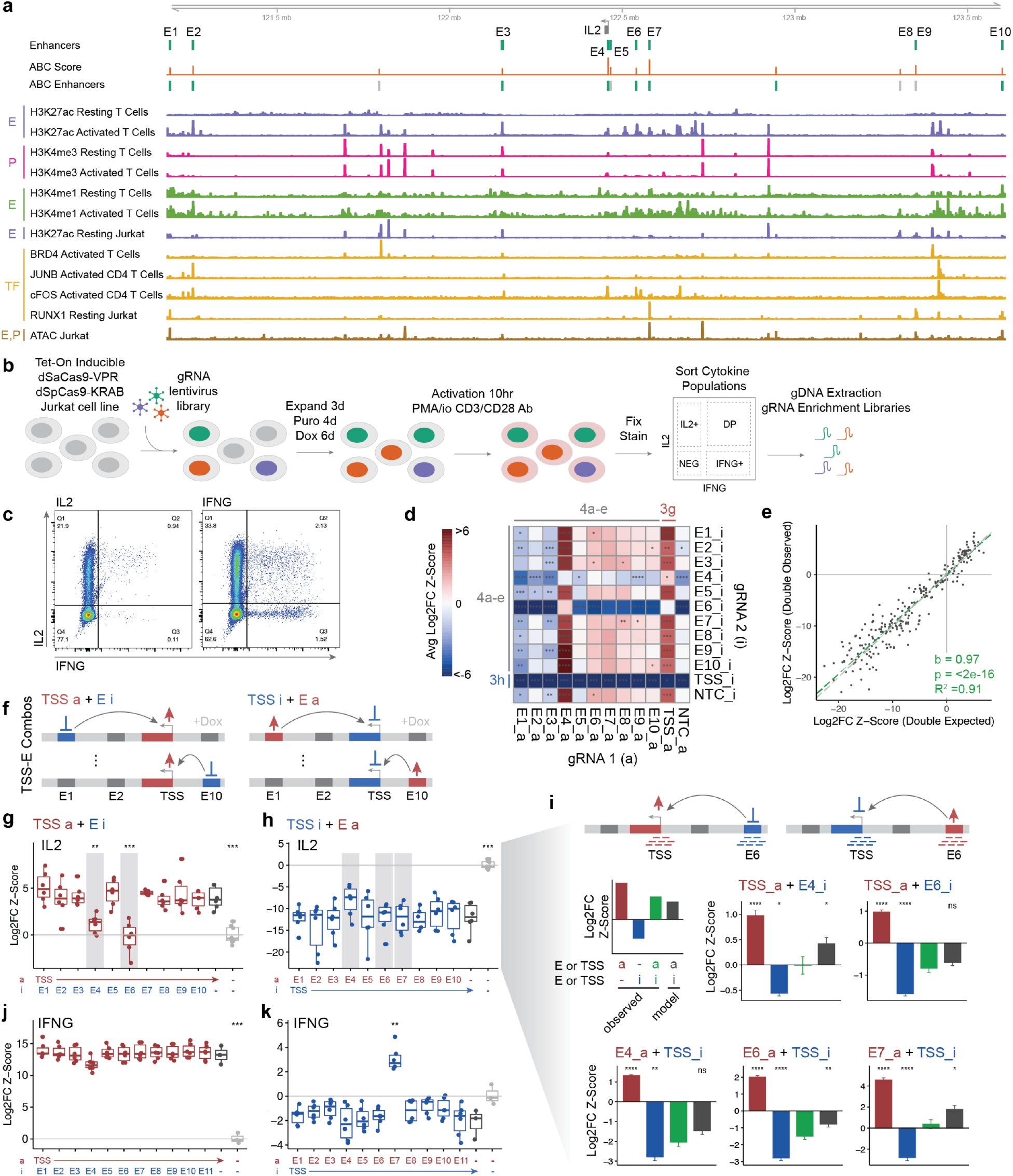
CRISPRai defines hierarchies in transcriptional regulation between promoter and enhancers. (a) Genome tracks showing regulatory and epigenetic landscape of *IL2* gene locus, with tracks showing: enhancers included in screen; ABC Score and predicted *IL2* enhancers using data from resting (gray) and CD3/PMA activated (green) Jurkat T cells^42^; ChIP-seq: primary T cell H3K27ac, H3K4me3, and H3K4me1 in resting and activated cells^64,65^, resting Jurkat H3K27ac^38^, activated primary T cell BRD4^84^, activated primary CD4 T cell JUNB and cFOS^80^, resting Jurkat RUNX1^38^; resting Jurkat ATAC-seq^85^. (b) Schematic of CRISPRai screen to investigate regulatory landscape of cytokine genes in Jurkat T cells. (c) Representative intracellular staining of cytokines IL2 and IFNG upon activation in Jurkat T cells. (d) Heatmap showing average log2FC gRNA enrichment (IL2+ / NEG) of all single, double, and NTC gRNA pairs included in the screen. 2 gRNAs per enhancer (2 a, 2 i). (e) Scatter plot of log2FC gRNA enrichment (IL2+ / NEG) for all TSS-E doubles, showing expected and observed gRNA enrichment. Expected enrichment calculated using additive model of singles. Linear regression slope (b), p value (p), and correlation coefficient (R^2^) are shown. (f) Schematic of pairwise combinations of bidirectional promoter-enhancer perturbations (TSS-E combinations). (g) TSS hierarchy over enhancers for *IL2* gene for TSS-CRISPRa and enhancer-CRISPRi pairs. Log2FC gRNA enrichment (IL2+ / NEG) is shown. (h) Same as g, for TSS-CRISPRi and enhancer-CRISPRa pairs. (i) Examples of specific TSS-E combinations highlighted in g,h with gray bars. (Top) schematic of validation screen setup with each enhancer tiled by 8 gRNAs. (Middle, bottom) plots show log2FC gRNA enrichment (IL2+ / IL2-). Bins represent singles (a or i), double (ai), and expected double from additive model (ai model). (j) Same as g for *IFNG* gene. Log2FC gRNA enrichment (IFNG+/ NEG) is shown. (k) Same as h for *IFNG* gene. Log2FC gRNA enrichment (IFNG+ / NEG) is shown. All data from *IL2* locus primary screen unless noted otherwise, n=3 biological screen replicates, 2 gRNAs per enhancer. j,k *IFNG* locus screen data, n=3 biological screen replicates, 2 gRNAs per enhancer. i *IL2* locus validation screen data from n=3 biological replicates, 8 gRNAs per enhancer, data displayed for 2-8 gRNAs. Significance tested relative to (g,h,j,k) TSS single perturbation and (i) observed double perturbation. d,i p values are calculated by Wilcoxon test. g,h,j,k p values are calculated by t test. Significance cutoffs: ns p > 0.05, * p <= 0.05, ** p <= 0.01, *** p <= 0.001, **** p <= 0.0001.

After identifying that a subset of enhancers was capable of counteracting TSS perturbation, we investigated these enhancers further. We designed a second gRNA pool to validate findings from the first screen and investigate enhancer function over a broader genomic range. We selected a subset of enhancers from the first *IL2* locus screen and tiled each enhancer with 8 gRNAs (**Fig. 3i top, Supplementary Fig. S9g**), including all E-E and TSS-E CRISPRai pairings as well as NTCs, for a pool of 4,096 gRNA pairs (3,136 doubles, 896 singles, and 64 NTCs, **Supplementary Table S5**), and constructed gRNA enrichment libraries for IL2+ and IL2-populations (**Supplementary Fig. S9a-h**). Enhancers were selected for inclusion in the validation screen based on their demonstrated potential for a strong functional effect or a genetic interaction from the first screen. Log2FC gRNA enrichment z-scores ranged from ∼ −5 with CRISPRi to ∼ +7.5 with CRISPRa in the validation screen (**Supplementary Fig. S9e**), and true gRNA enrichment FC ranged from 0.75 to 1.35 (not shown). E2_i_5_val and E9_i_2_val had some off targets within 2kb of the TSS, however these were low risk (mismatch=4, bulge size=1) (not shown) and did not appear to affect gRNA behavior as determined by their concordance with other gRNAs for the corresponding enhancer that had zero TSS proximal off-targets (Cas OFFinder, parameters: mismatches <=4, DNA and RNA bulge size <=1)^78^ (**Supplementary Fig. S9h**). The validation screen confirmed the gatekeeper effects of E4 and E6 in the TSS-E interactions and highlighted the presence of strong functional hotspots within other enhancers that were also capable of counteracting TSS perturbation (**Fig. 3i**). When comparing true IL2+ gRNA enrichment FC in single perturbations, E4 and E6 exhibited 24% and 66% of TSS repression and 0.932 and 0.814 FC relative to NTC (TSS FC 0.719) with CRISPRi, and E4, E6, and E7 activation exhibited 104%, 115%, and 160% of TSS activation and 1.20, 1.33, and 1.85 FC relative to NTC (TSS FC 1.15) with CRISPRa, respectively (not shown). Across the E6 enhancer, E6 repression was capable of strongly counteracting TSS activation by switching the target gene into the off state; log2FC gRNA enrichment z-score was negative indicating that this gRNA pair was strongly depleted in IL2+ cells, and thus prevented *IL2* expression (**Fig. 3i middle, Supplementary Fig. S9h**). The effect for E4 was weaker and varied across the enhancer, with log2FC gRNA enrichment z-score of ∼0, equal to the enrichment of NTC gRNAs. The validation screen demonstrated similar E4 and E6 behavior in the reverse condition, albeit to a lesser extent since overall TSS repression was stronger than TSS activation (**Fig. 3i bottom**). Additionally, we found a hotspot in E7 whose activation strongly counteracted TSS repression (**Fig. 3i bottom**). We noted that CRISPRa appears to be more focal than CRISPRi, possibly due to different mechanisms of chromatin remodeling induced by VPR and KRAB, respectively (**Supplementary Fig. S7h, Supplementary Fig. S9h**).

### Hierarchies in enhancer-enhancer interactions for target gene control

We next investigated how enhancers act in concert, or interact with other enhancers, to exert control over the target gene. We compared the log2FC gRNA enrichment z-scores of various E-E doubles from the *IL2* locus validation screen (**Fig. 4a**). Similar to the TSS-E doubles, E-E doubles largely followed an additive model with respect to log2FC gRNA enrichment z-score (R^2^=0.75, b=0.95, p=<2e-16, **Fig. 4b**). A Fisher’s Exact test on the validation gRNA pool for enrichment of specific enhancers in the tails of the residual distribution confirmed the general adherence to the additive model (**Supplementary Fig. S9j**). We noted that single and double E-E perturbations enabled tuning of *IL2* expression over a broad range, supporting a hypothesis that multiple enhancers of varying strengths enable more precise tuning collectively than would be possible with fewer enhancers (**Fig. 4c**). Importantly, we found that the gatekeeper enhancers identified from the TSS-E double perturbations, E4 and E6, showed similar gatekeeper behavior when paired with other enhancers (**Supplementary Fig. S9i**). Double perturbations that contained activation of E4 or E6 were enriched in IL2+ cells compared to the merged set of doubles containing all other enhancer pairings, while gRNAs for double perturbations in which E4 and E6 were repressed were depleted from IL2+ cells (**Supplementary Fig. S9i**). In other words, activation of E4 or E6 drove gene expression when other enhancers in the same locus were silenced, and conversely, silencing E4 or E6 prevented gene expression even if other *IL2* enhancers were activated.

**Fig. 4.**
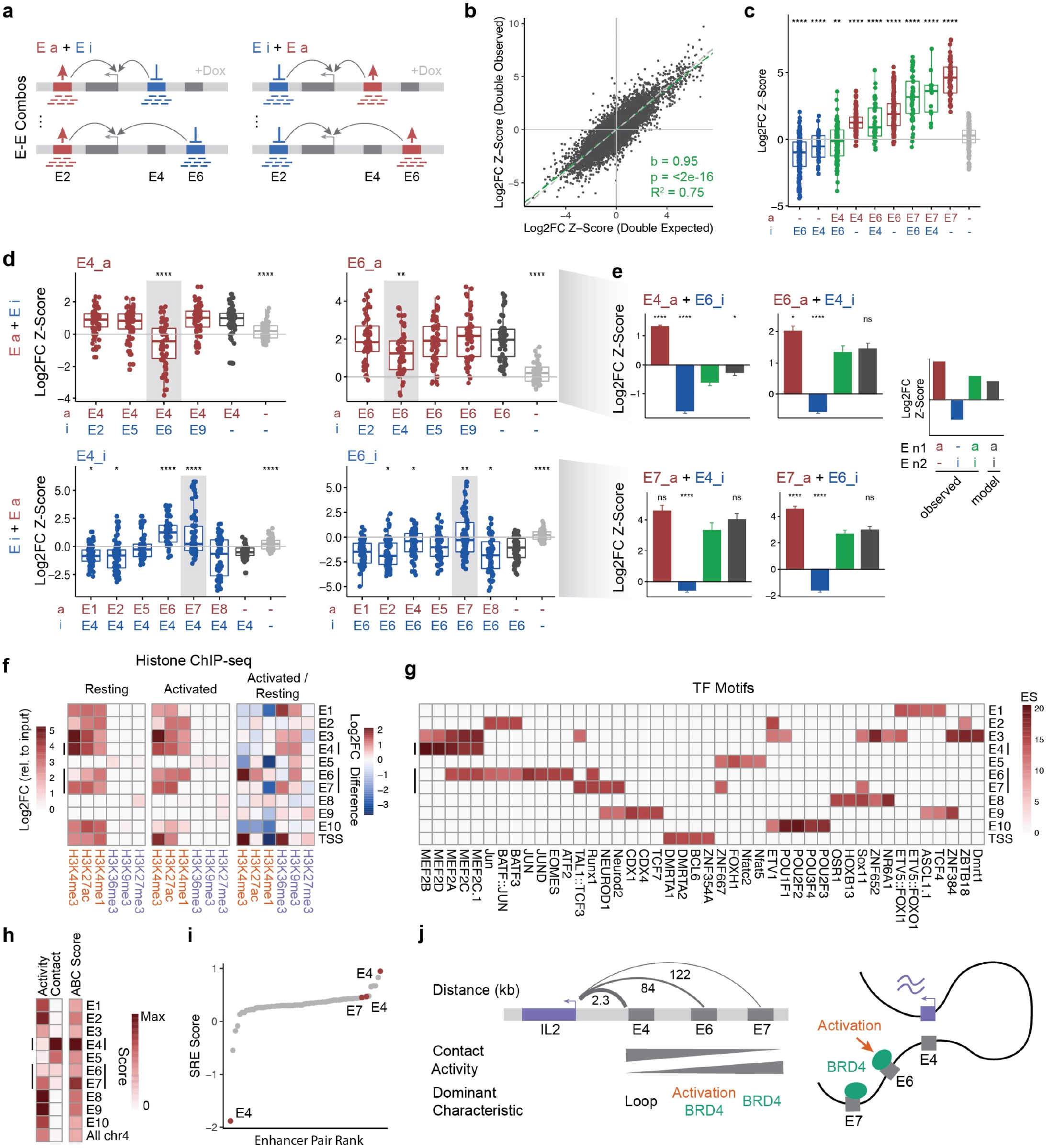
Hierarchies in enhancer-enhancer interactions for target gene control. (a) Schematic of pairwise combinations of E-E perturbations (E-E combinations) and tiling of enhancers with 8 gRNAs in *IL2* locus validation screen. (b) Scatter plot of log2FC gRNA enrichment (IL2+ / IL2-) for all E-E doubles, showing expected and observed log2FC gRNA enrichment. Expected enrichment calculated using additive model of singles. Linear regression slope (b), p value (p), and correlation coefficient (R^2^) are shown. (c) Log2FC gRNA enrichment (IL2+ / IL2-) for selected singles and doubles for E-E pairs showing tuning of target gene expression. Each pair of perturbed enhancers is indicated. (d) Log2FC gRNA enrichment (IL2+ / IL2-) for all E-E pairs containing (left) E4 and (right) E6 for (top) activation or (bottom) repression of E4 and E6, respectively. (e) Examples of CRISPRai for specific E-E combinations containing E4, E6, and a hotspot in E7. Bins represent singles (a or i), double (ai), and expected double from additive model (ai model). Data displayed for 2-8 gRNAs. (f) ENCODE histone ChIP-Seq^64,65^ from resting and activated primary T cells. Peaks overlapping each enhancer are aggregated. Activating marks (orange) and repressive marks (purple) are shown. (g) JASPAR TF motif^79^ enrichment in each enhancer relative to other enhancers. Lowercase denotes variant motif. (h) Activity, contact, and ABC scores from the ABC model^42^ for each enhancer. (i) SRE scores for all E-E pairings in SRE model^36^. Selected enhancer pairs with high magnitude SRE scores that also contain enhancers from our screen are annotated with the CRISPRai enhancer in the pair. (j) Proposed model of enhancer-mediated gene regulation for *IL2* by strong functional enhancers. ABC model contact and activity scores from h are summarized. Data from *IL2* locus validation screen, n=3 biological replicates, 8 gRNAs per enhancer. P values are calculated from Wilcoxon test. Significance tested relative to (d) single perturbation of indicated enhancer and (e) observed double perturbation. Significance cutoffs: ns p > 0.05, * p <= 0.05, ** p <= 0.01, *** p <= 0.001, **** p <= 0.0001.

To investigate this behavior in more detail, we looked at the interaction between E4 and E6 with each individual other enhancer included in the pool. E6 repression counteracted E4 activation (**Fig. 4d, top left, bottom right**), and conversely, E6 activation counteracted E4 repression (**Fig. 4d bottom left, top right**), supporting the strong and moderate gatekeeper roles of E6 and E4, respectively. E7 activation was also capable of counteracting E4 repression (**Fig. 4d bottom left**), similar to what we observed with the TSS (**Fig. 3g-i**). All other enhancers had minimal to no ability to counteract E4 and E6 perturbation (**Fig. 4d**). Interestingly, E1 and E2 activation weakly reduced log2FC gRNA enrichment, suggesting that these two enhancers, which are both ∼1.2 Mb from the TSS, may be weak repressive regulatory elements (**Fig. 4d, Supplementary Fig. S9h**). Examples of selected E-E double perturbations are shown, along with the expected double log2FC gRNA enrichment under an additive model calculated from the corresponding singles (**Fig. 4e**). Bidirectional epigenic editing of E4 and E6 enables reversible control of target gene expression as indicated by opposite magnitude gRNA enrichment in the ai versus ia conditions, and the interaction between E4 and E6 was additive regardless of the perturbation direction (ai vs ia) (**Fig. 4e top**). E7 activation strongly counteracted both E4 and E6 repression due to the large magnitude of E7 gRNA enrichment in IL2+ cells (**Fig. 4e bottom**). Each of these E-E pairs showed additive or near-additive behavior (**Fig. 4e**).

We then investigated the potential molecular mechanism to explain the gatekeeper function of E4 and E6, and the strong activator function of E7. We hypothesized that these strong functional enhancers may have unique histone marks or TF motif enrichment signatures compared to other enhancers. First, we compared ENCODE histone ChIP-seq in resting and activated primary T cells^64,65^ (**Fig. 4f, Supplementary Fig. S10a-d**), a relevant comparison since our screen endpoint was T cell activation (**Fig. 3b**). We found that E4 and E7 had high to moderate activating histone marks, including H3K4me3 and H3K27ac (**Fig. 4f, Supplementary Fig. S10b,d**). In contrast, E6 was relatively low for these histone marks, but showed a large increase in activating histone marks in activated compared to resting cells (**Fig. 4f**). The most prominent histone mark on E6 was H3K4me1, which is often enriched at primed enhancers (**Fig. 4f, Supplementary Fig. S10c**). This suggests that E6 is a primed and highly activation-responsive enhancer, which may contribute to its strong gatekeeper function, and that this function may be relevant in both the cell line and primary T cell context (**Fig. 4f**). In addition, compared to other enhancers, E6 was highly enriched in JASPAR TF motifs^79^ involved in T cell activation, including BATF3, JUN, JUND, ATF2, and EOMES, further supporting the activation-responsive nature of E6 and its potential importance in regulating activation-induced *IL2* expression in T cells (**Fig. 4g**). AP-1 family TF occupancy at E6 was corroborated with ChIP-seq in activated primary CD4 T cells (**Supplementary Fig. S10c**)^80^.

Additionally, we investigated the enhancer characteristics derived from the data in the ABC Model for gene-specific enhancer prediction^41,42^. Under the ABC Model, E4 has the highest predicted enhancer score of all the enhancers, with high contact score (high contact frequency with the TSS, mixed cell type Hi-C), yet low activity score in (combined score of ATAC-seq and H3K27ac ChIP-seq, Jurkat T cells)^42^ (**Fig. 4h**). Thus, E4 gatekeeper function is likely largely driven by frequent E-P contacts rather than enhancer activity. Conversely, E7 had the opposite situation, with low contact score, but high activity score, resulting in a relatively high overall predicted enhancer score (**Fig. 4h**). Taken together, these attributes of E4 and E7 indicate that a complementary mechanism may exist between E-P contacts and enhancer activity, which enable either property to drive enhancer-mediated gene regulation in different contexts. Further, Lin et al. reported the finding that while some enhancers exhibit additive behavior with other enhancers, certain enhancers, termed synergistic regulatory enhancers (SREs), exhibit synergistic E-E interactions, that these interactions usually occur between distal enhancers, and that they overlap with disease variants^36^. We investigated whether any strong functional CRISPRai enhancers overlapped with the SRE enhancer set identified in Lin et al. We found that E7 and most notably E4, were present in the top most synergistic SRE E-E pairs, confirming their importance in *IL2* gene regulation (**Fig. 4i**). E6 was not present in the SRE enhancer set. We summarize a proposed model of gene regulation for the strong functional enhancers identified by CRISPRai, highlighting that different enhancers likely function through different regulatory modes (**Fig. 4j**).

## DISCUSSION

We developed a bidirectional epigenetic editing system, termed CRISPRai, leveraging CRISPRa and CRISPRi to expand the toolkit for investigating genetic interactions and noncoding genetic elements. We demonstrated the utility of the system in perturbing pairs of genes with a single-cell RNA-seq readout and applied this to study the context-specific genetic interaction between SPI1 and GATA1, two key hemopoietic lineage TFs. Further, we applied CRISPRai to investigate how multiple enhancers act in concert to control target gene expression, which revealed 1) that the promoter holds hierarchy over enhancers for target gene regulation, 2) that pairwise enhancer perturbations enable fine tuning of target gene expression, and 3) the existence of strong functional gatekeeper enhancers with the power to turn off target genes despite promoter activation.

Enhancer functionality is heterogenous and molecularly complex. Some enhancers may act in an additive manner^76^, while other rare enhancers may have synergistic effects in combination^36^. Some enhancers offer redundancy, while others are dominant levers for gene expression control^76,77,81,82^. Enhancers differ in their structural chromatin contacts^83^, chromatin modifications^39^, and in which TFs they are capable of recruiting^43^, which likely governs their function and the target genes for which they are compatible. These characteristics are consistent with our findings that combined enhancer function is a primarily additive and with the presence of dominant gatekeeper enhancers, a novel finding from this study.

It is known that individual enhancers modulate target gene expression, typically in an activating manner, and that this function is at least in part related to E-P distance, chromatin modifications, and enhancer-specific TF recruitment^37,39,44^, but how different enhancers function in concert to control a target gene or why some genes have more enhancers than others is unknown. We found that applying different single and double perturbations of different enhancer combinations using bidirectional epigenetic editing by CRISPRai enabled tuning of gRNA enrichment in IL2+ versus IL2-cells over a range of values (**Fig. 4c**). This tuning depending on the directionality and combination of enhancer perturbation suggests that enhancers have complex interactions whose purpose may be to collectively enable precise tuning of target gene expression. Rather than having discrete levels of target gene activation, input from multiple enhancers may enable cells to modulate target gene expression with much finer control than would be possible with fewer enhancers. Further, our data suggests that the underlying mechanisms that govern enhancer function are complex and variable across different enhancers, since we found different histone modification, TF motif, activity, and chromatin contact signatures across strong functional enhancers (**Fig. 4f-h,j**).

Further application of CRISPRai to additional genomic loci will help elucidate general rules and hierarchies that govern gene regulation across different genomic regions. Our study of the *IL2* locus revealed a novel hierarchical relationship between enhancers and the promoter, and we anticipate that similar studies of other regions will elucidate regulatory mechanisms across the genome. Notably, enhancer complexity suggests that higher-order combinatorial interactions (e.g. 3+ loci), beyond pairwise (2 loci) perturbations performed here, may reveal further nuances of enhancer-mediated gene regulation. Additionally, greater combinatorial perturbations will likely reveal novel biological pathway insights. Further methods to broaden the toolkit for studying genetic interactions and noncoding elements, such as incorporating other types of epigenetic effector domains for combinational CRISPR perturbations and coupling with high-content readouts such as single-cell multiomics, will further enhance our understanding of the multifaceted and heterogenous mechanisms underlying gene regulation.

## SUPPLEMENTARY FIGURE LEGENDS (Supplementary Fig. S1-S10)

**Supplementary Fig. S1. (related to Fig. 1,3).**
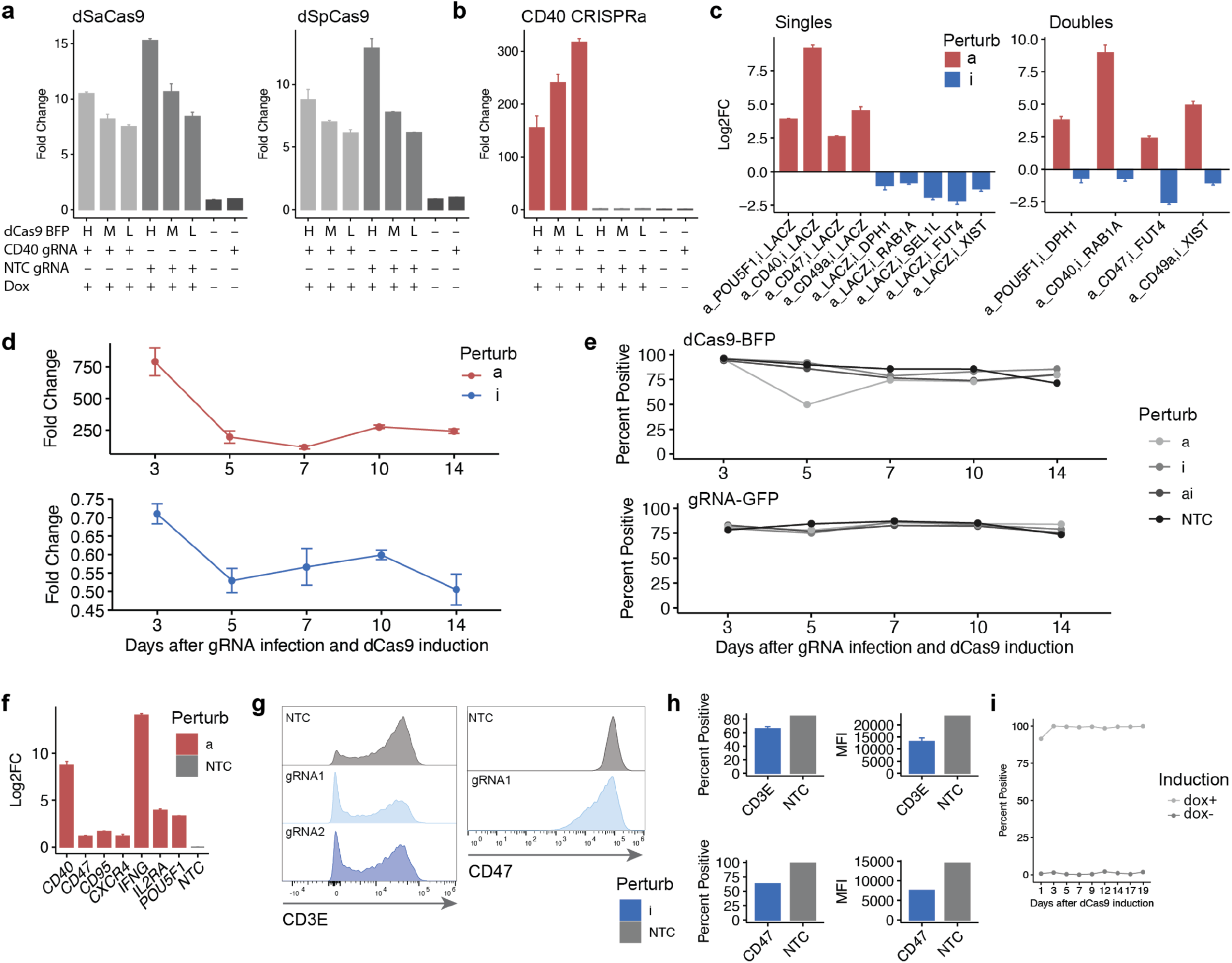
Development of CRISPRai system. (a-e) Development of K562 CRISPRai cell line in bulk (related to **Fig. 1**). (a,b) Demonstration of construct expression, inducibility of CRISPRai system and dependence on dCas9 cassette expression level (BFP). Expression level by qPCR of (a) dSaCas9 and dSpCas9, and (b) CD40 target gene. Data shows construct or gene expression after sorting high, mid, low (H,M,L) dCas9 (BFP) expression levels by FACS of cells expressing gRNAs for CD40 CRISPRa or NTC. (c) Log2FC gene expression by qPCR of several genes relative to NTC in single and double perturbations, highlighting bidirectional control. (d) Time course of gene expression during CRISPRai perturbation of 2 genes as single perturbations for (top) CRISPRa and (bottom) CRISPRi. Perturbation effect on target genes remains at least until day 14. (e) Time course of construct expression by flow cytometry of (top) dCas9 cassette (BFP) and (bottom) gRNA (GFP). dCas9 remains expressed at high levels for greater than 14 days and is not silenced. (f-i) Development of Jurkat CRISPRai cell line in bulk (related to **Fig. 3**). (f) Log2FC gene expression by qPCR of several genes relative to NTC in single perturbations for CRISPRa. (g) Protein expression by flow cytometry showing CRISPRi of 2 genes (CD3E and CD47). (h) Same data as g, showing percent positive and MFI. (i) Time course of dox induction showing dCas9 cassette (BFP) does not silence over a 20 day period. a-d,f all qPCR analysis was done using the double delta Ct method relative to housekeeping gene *ACTB* and NTC gRNAs. NTC gRNA is against *lacZ* gene.

**Supplementary Fig. S2. (related to Fig. 1-4).**
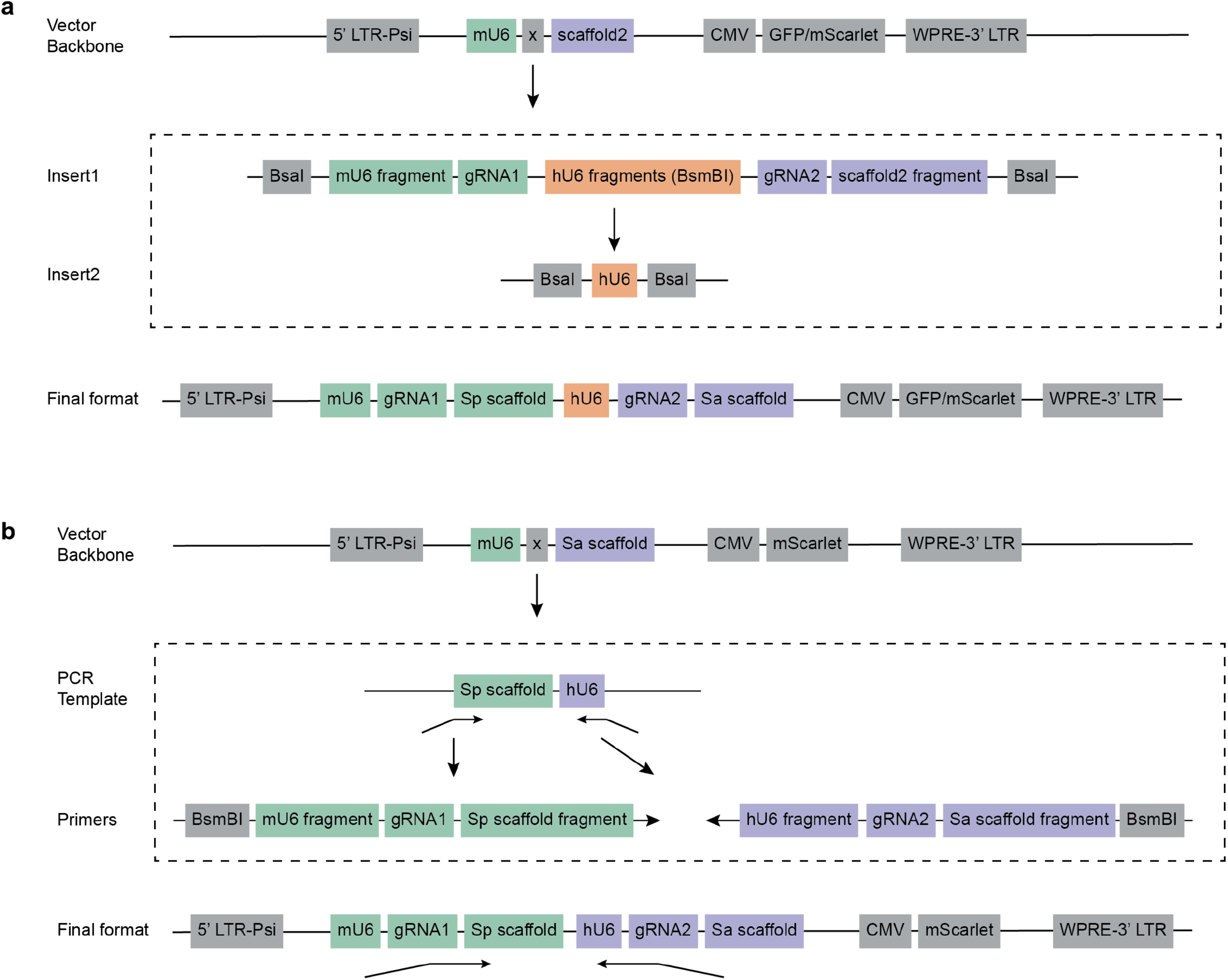
Dual CRISPR gRNA cloning. (a) Cloning strategy for dual gRNAs for K562 single-cell screen. (b) Cloning strategy for dual gRNAs for Jurkat gRNA enrichment screen.

**Supplementary Fig. S3. (related to Fig. 2).**
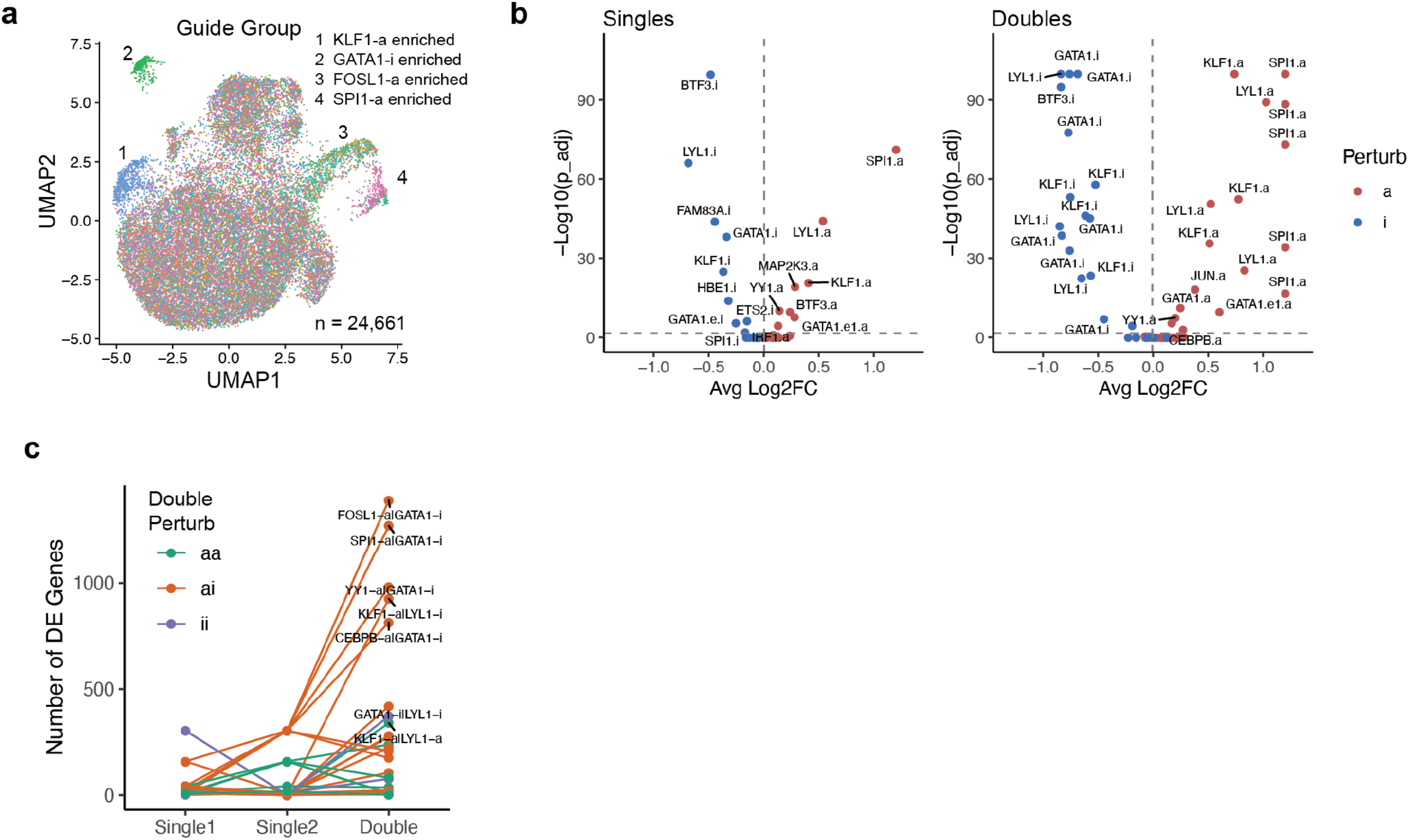
Additional analysis of single-cell CRISPRai screen in K562 cells. Data from single-cell CRISPRai K562 screen from **Fig. 1**,**2**. (a) UMAP representation of all single-cell transcriptomes from screen colored by gRNA group. Labeled clusters highlight cells with strong perturbation-driven phenotypes. (b) Volcano plots showing DE of CRISPRai target genes in (left) singles and (right) doubles. Horizontal dotted line at p_adj <= 0.05. (c) Data from all perturbation sets from K562 single-cell screen, showing number of DE genes in single and double perturbations for all perturbation sets with n > 50 cells per group. All cells, including non-perturbation driven cells, are included for DE testing. c significance cutoffs for DE genes are abs(log2FC) > 0.1, p_adj < 0.05.

**Supplementary Fig. S4. (related to Fig. 2).**
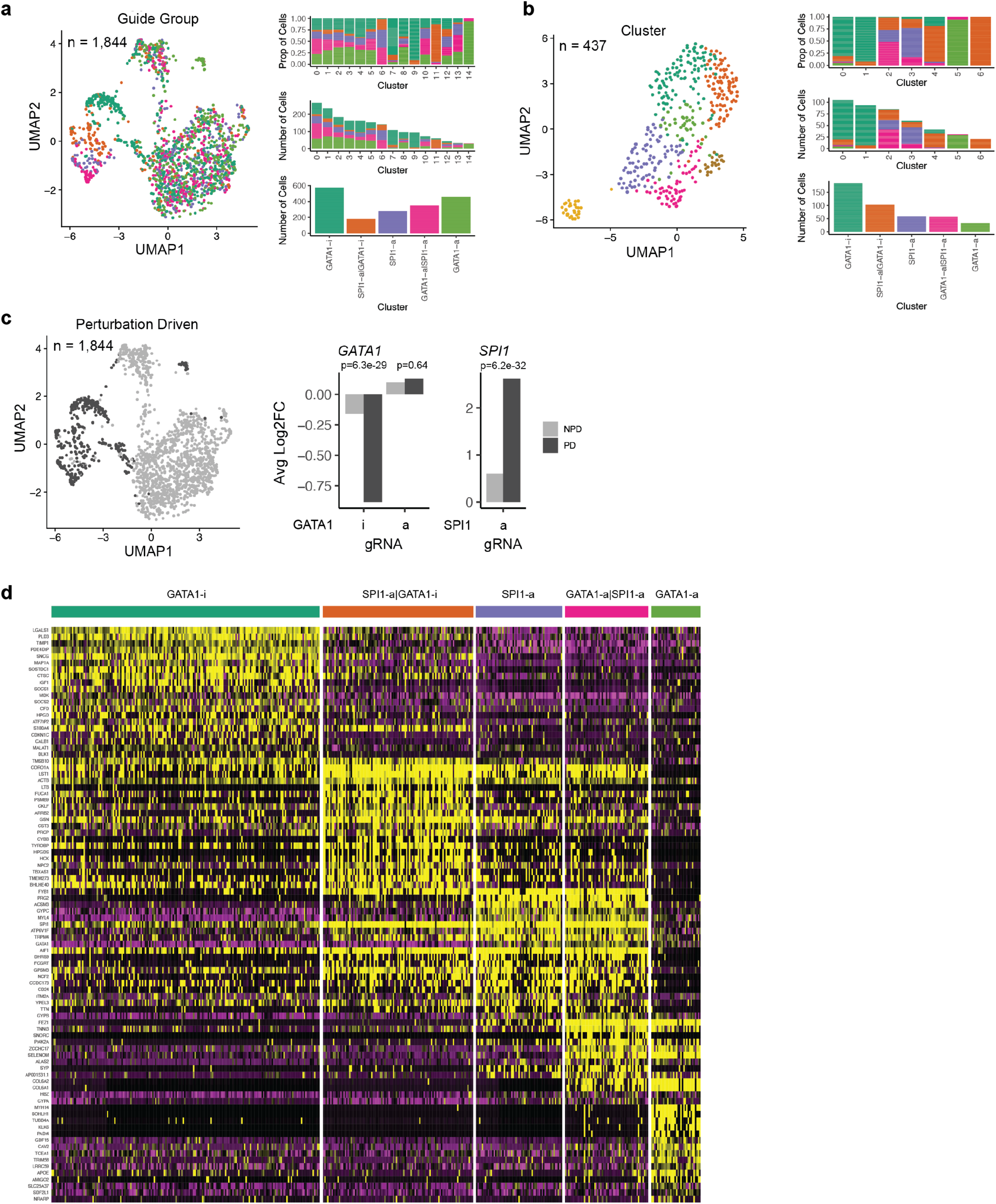
Quality control of SPI1 and GATA1 genetic interaction. Data from single-cell CRISPRai K562 screen from **Fig. 1,2**. (a) UMAP subclustering of all cells with gRNA calls in the SPI1-GATA1 perturbation set. Proportion or number of cells per gRNA group is shown. (b) Same as a for perturbation-driven cell subset. (c) (Left) Same data as a, colored by whether cells had perturbation-driven (PD) or non-perturbation driven (NPD) clustering. Perturbation driven cells were defined as cells in all clusters that do not comprise equal proportions of cells from each gRNA group. (Right) Average log2FC gene expression of *GATA1* and *SPI1* in PD and NPD cells. (d) Heatmap of top DE genes by gRNA group. c,d DE gene testing for each gRNA group is against NTC. d significance cutoffs for DE genes are abs(log2FC) > 0.25, p_adj < 0.05.

**Supplementary Fig. S5. (related to Fig. 2).**
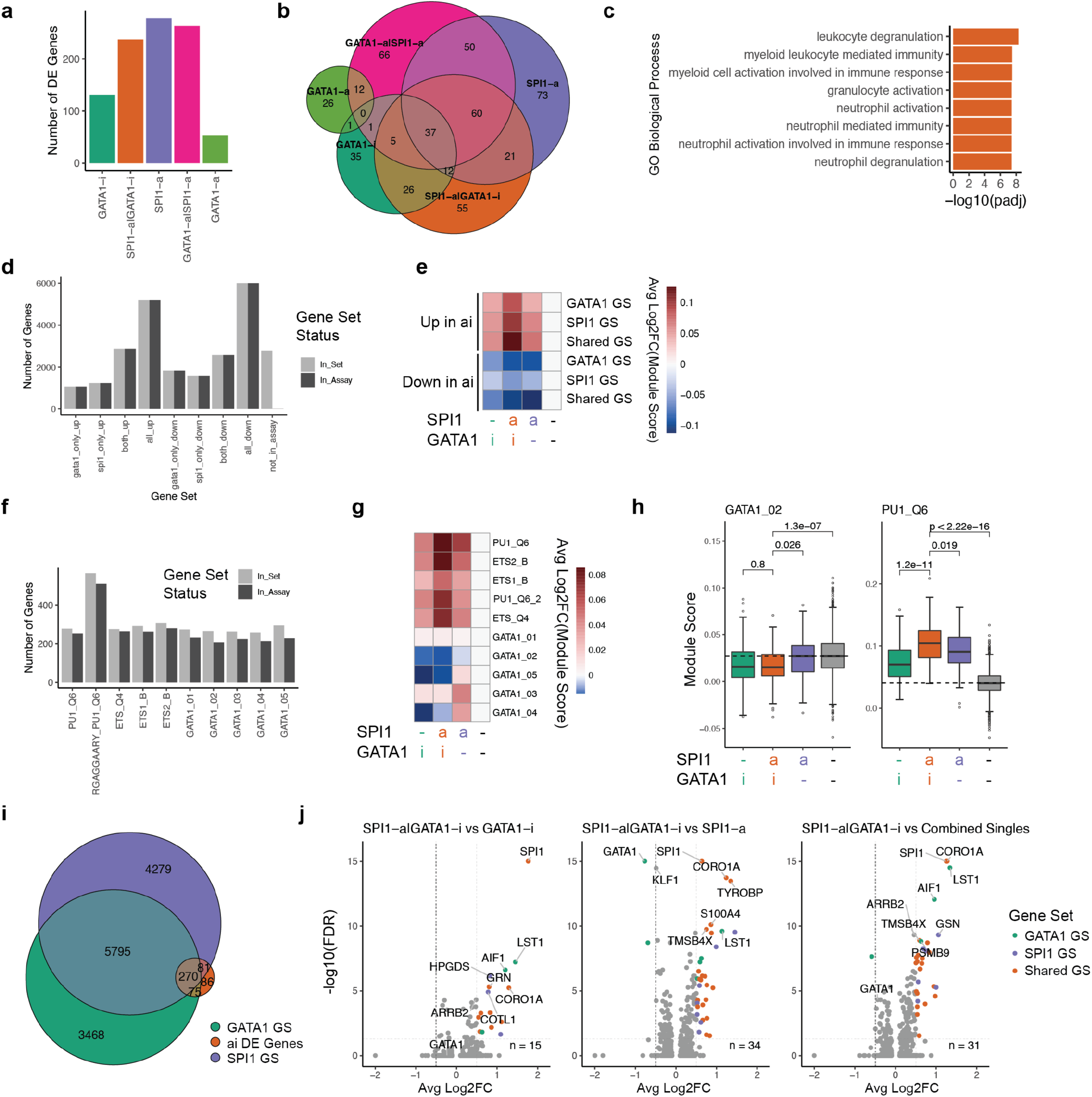
Additional analysis for SPI1 and GATA1 genetic interaction. Data from single-cell CRISPRai K562 screen from **Fig. 1,2**. (a) Number of DE genes by gRNA group. (b) Overlap of DE genes across gRNA groups. (c) GO biological process term enrichment for upregulated genes uniquely DE in SPI1-a|GATA1-i double perturbation (a subset of 58 genes from venn diagram in **Fig. 2c**). Genes are enriched for lineage-and target-specific processes. (d) Number of genes in each ENCODE TF target gene set^64–66^ and the subset that is present in the dataset for GATA1 only, SPI1 only, or shared TF target genes partitioned by up or down regulation in the SPI1-a|GATA1-i double perturbation. All_up and all_down gene sets are shown in **Fig. 2f,g**. (e) Average log2FC (relative to NTC) gene expression module score for gene sets in d by gRNA group. Up or down regulation in double perturbation (SPI1-a|GATA1-i) is indicated. (f) Number of genes in each MSigDb TF target gene set^67,68^ and the subsets that are present in the dataset for SPI1 (PU.1), GATA1, or ETS family gene sets. (g) Average log2FC (relative to NTC) gene expression module score for gene sets in f by gRNA group. (h) Examples from g. (i) Overlap between statistically significant upregulated DE genes in the SPI1-a|GATA1-i double perturbation and SPI1 and GATA1 ENCODE TF target gene sets from d. (j) DE genes in SPI1-a|GATA1-i double perturbation relative to individual single or combined single perturbations. Colored by membership in ENCODE TF target gene set. Only genes in ENCODE TF target genes sets were tested for DE. Significance cutoffs for DE genes are (a-c,j) abs(log2FC) > 0.5, p_adj < 0.05, and (c,i) log2FC > 0.5, p_adj < 0.05. DE gene testing for each gRNA group is against NTC, all detected genes considered unless noted otherwise.

**Supplementary Fig. S6. (related to Fig. 3).**
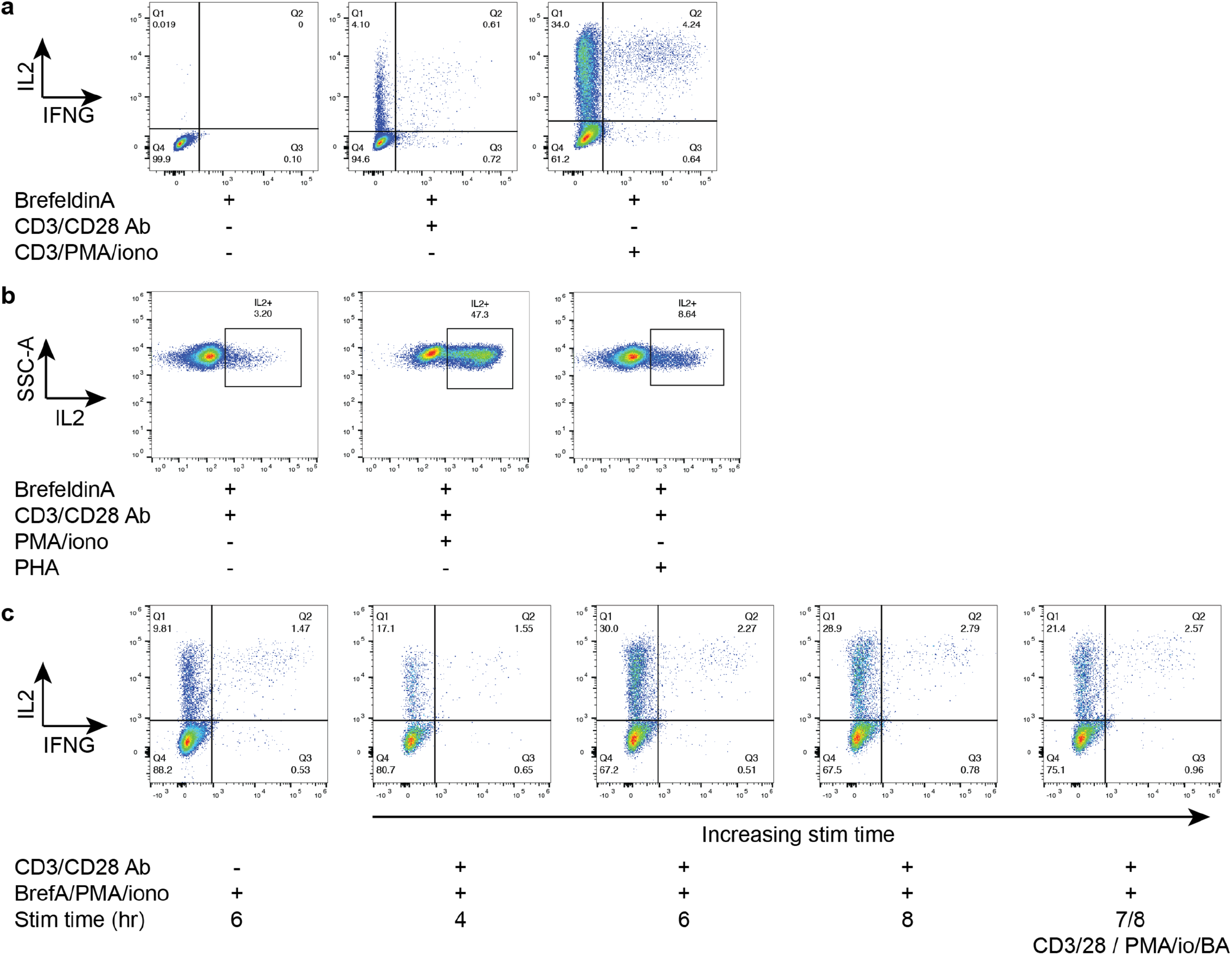
Optimization of Jurkat T cell activation. Flow cytometry of intracellular cytokine staining for IL2 and IFNG in Jurkat T cells in various conditions. (a) Comparison of CD3/CD28 and PMA/ionomycin for 10hr activation. (b) Comparison of combinations of CD3/CD28, PMA/ionomycin, and PHA for 10hr activation. (c) Time course of CD3/CD28 and PMA/ionomycin activation.

**Supplementary Fig. S7. (related to Fig. 3).**
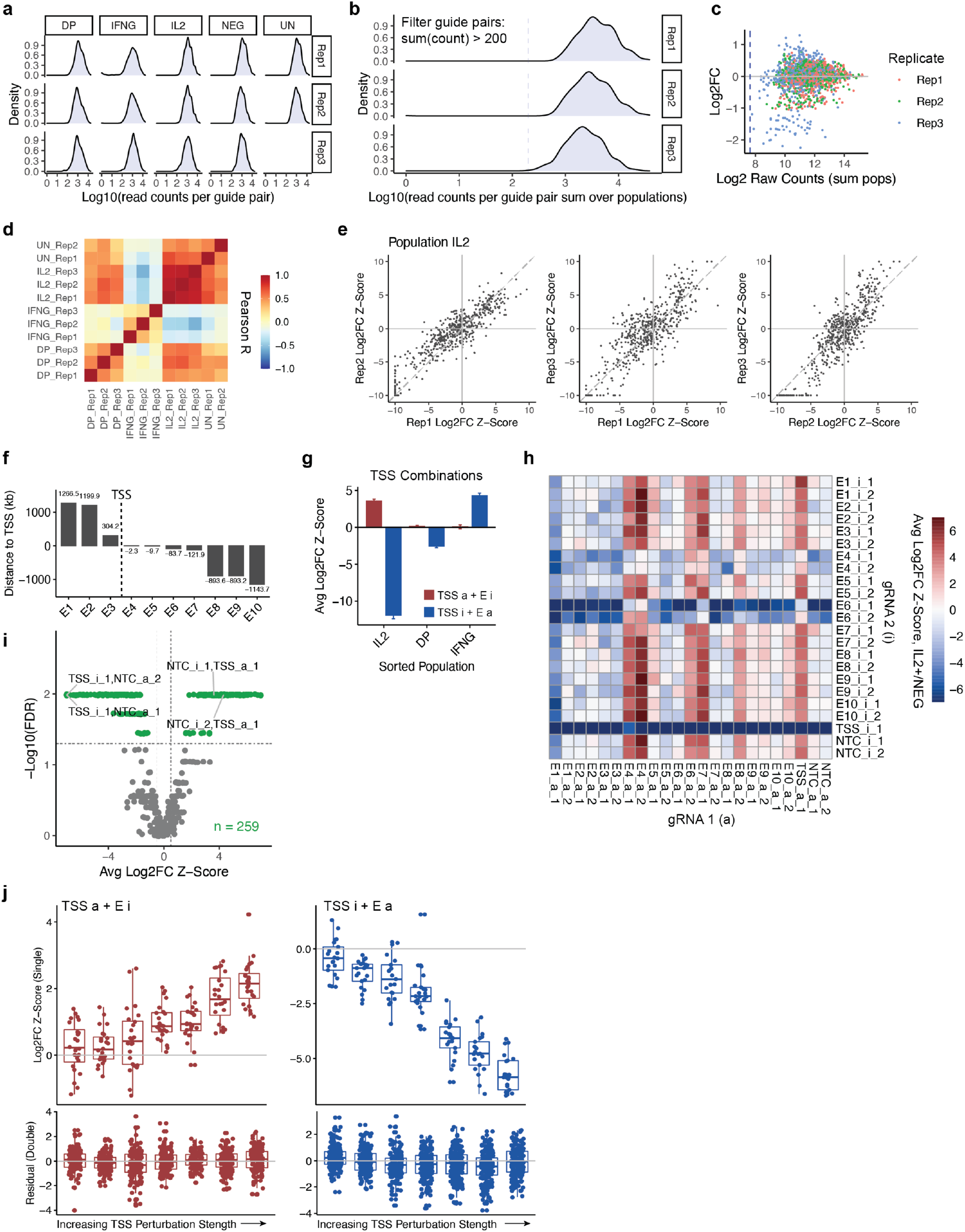
Additional data from Jurkat *IL2* locus primary screen and TSS-E perturbations. (a) Log10 raw read counts per gRNA pair by sorted population and biological replicate. DP=IL2+IFNG+ (double positive), IFNG=IL2-IFNG+, IL2=IL2+IFNG-, NEG=IL2-IFNG-(double negative), UN=unsorted. (b) Log10 raw read counts per gRNA pair summed over all populations, with read count cutoff (blue dotted line) used for filtering reads (filter: sum(count) > 200). (c) Log2FC gRNA enrichment versus log2 raw read counts per gRNA, colored by biological replicate. (d) Pearson correlation of log2FC gRNA enrichment (IL2+ / NEG) across all gRNAs for sorted populations and biological replicates. (e) Log2FC gRNA enrichment correlation of biological replicates for IL2 sorted population. (f) Distance between enhancer midpoint and *IL2* TSS. (g) Average log2FC gRNA enrichment (IL2+ / NEG) for all single and double TSS-E combinations, binned by sorted population. (h) Overview of screen data showing all singles, doubles, and NTCs included in the screen. Columns and rows indicate gRNA1 and gRNA2 in the pair. Average log2FC gRNA enrichment (IL2+ / NEG) is shown. 2 gRNAs per enhancer. (i) Volcano plot showing statistically significant log2FC gRNA enrichment (IL2+ / NEG) for all gRNA pairs. Includes all singles, doubles, and NTCs. TSS singles are highlighted. Significance cutoff FDR < 0.05, log2FC z-score > 0.5. (j) Data from *IL2* validation screen. Additive behavior in TSS-E doubles holds across range of TSS perturbation strengths, for TSS activation (left) and TSS repression (right). (Top) log2FC gRNA enrichment (IL2+ / IL2-) of TSS gRNAs in singles. (Bottom) residual of each gRNA double from the additive model linear fit, binned by perturbation strength of the TSS gRNA in the pair. a-i data from *IL2* primary screen, n=3 biological replicates, 2 gRNAs per enhancer. j data from *IL2* validation screen, n=3 biological replicates, 8 gRNAs per enhancer.

**Supplementary Fig. S8. (related to Fig. 3).**
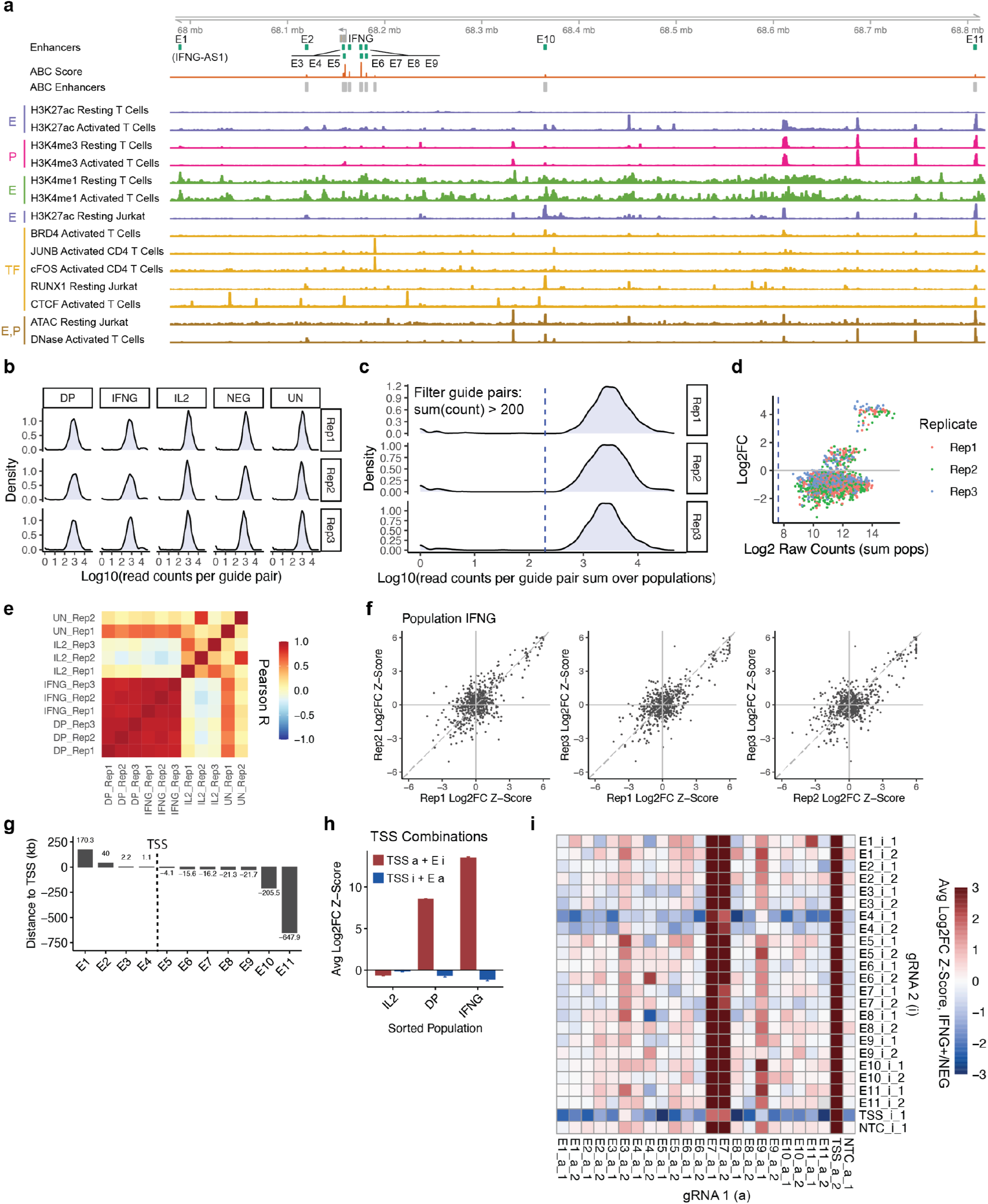
Additional data from Jurkat *IFNG* locus primary screen. (a) Genome tracks showing regulatory and epigenetic landscape of *IFNG* gene locus. Same tracks as **Fig. 3a**, with additional tracks: activated T cell CTCF ChIP-seq and activated T cell DNase-seq^64,65^. (b) Log10 raw read counts per gRNA pair by sorted population and biological replicate. DP=IL2+IFNG+ (double positive), IFNG=IL2-IFNG+, IL2=IL2+IFNG-, NEG=IL2-IFNG-(double negative), UN=unsorted. (c) Log10 raw read counts per gRNA pair summed over all populations, with read count cutoff (blue dotted line) used for filtering reads (filter: sum(count) > 200). (d) Log2FC gRNA enrichment versus log2 raw read counts per gRNA, colored by biological replicate. (e) Pearson correlation of log2FC gRNA enrichment across all gRNAs for sorted populations and biological replicates. (f) Log2FC gRNA enrichment correlation of biological replicates for IFNG sorted population. (g) Distance between enhancer midpoint and *IFNG* TSS. (h) Average log2FC gRNA enrichment (IFNG+ / NEG) for all single and double TSS-E combinations, binned by sorted population. (i) Overview of screen data showing all singles, doubles, and NTCs included in the screen. Columns and rows indicate gRNA1 and gRNA2 in the pair. Average log2FC gRNA enrichment (IFNG+ / NEG) is shown. n=3 biological replicates, 2 gRNAs per enhancer.

**Supplementary Fig. S9. (related to Fig. 3,4).**
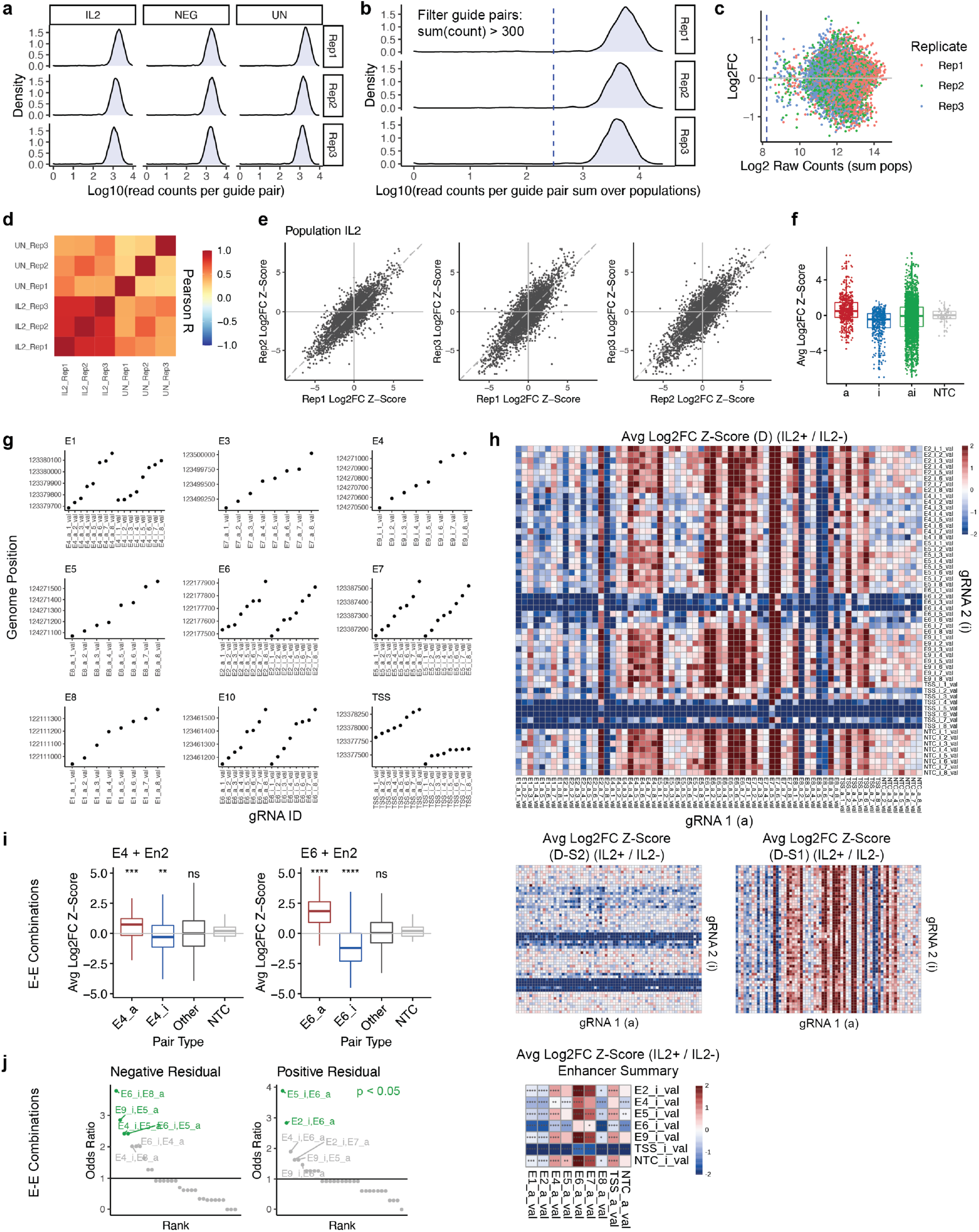
Additional data from Jurkat *IL2* locus validation screen. (a) Log10 raw read counts per gRNA pair by sorted population and biological replicate. IL2=IL2+, NEG=IL2-, UN=unsorted. (b) Log10 raw read counts per gRNA pair summed over all populations, with read count cutoff (blue dotted line) used for filtering reads (filter: sum(count) > 300). (c) Log2FC gRNA enrichment versus log2 raw read counts per gRNA, colored by biological replicate. (d) Pearson correlation of log2FC gRNA enrichment across all gRNAs for sorted populations and biological replicates. (e) Log2FC gRNA enrichment correlation of biological replicates for IL2 sorted population. (f) Average log2FC gRNA enrichment (IL2+ / IL2-) for all singles (a or i), doubles (ai), or NTCs included in the screen. (g) gRNA identity (gRNA ID) versus genome position. gRNAs are tiled across enhancers. gRNA ID number correlates with genome position. (h) Overview of screen data showing all singles, doubles, and NTCs included in the screen. (Top) Average log2FC gRNA enrichment as observed (D) (IL2+ / IL2-), (middle, left) with CRISPRa single subtracted (D-S2), and (middle, right) with CRISPRi single subtracted (D-S1). (Middle) Highlights contribution of non-subtracted single in the double gRNA pair condition. (Bottom) Average log2FC gRNA enrichment averaged over 8 gRNAs per enhancer. (i) Gatekeeper behavior of (left) E4 and (right) E6. Log2FC gRNA enrichment (IL2+ / IL2-) is shown. E-E doubles are grouped into 4 bins as follows: 1) all pairs with E4_a gRNA, 2) all pairs with E4_i gRNA, 3) all pairs without an E4 gRNA, 4) all NTCs. The same bins are also displayed for E6. (j) Fisher’s Exact test odds ratio for presence of E-E double gRNA pairs in the tails (top or bottom 5%) of the residual distribution of the additive model. Positive residual: observed double > expected double, negative residual: observed double < expected double. Statistical significance is highlighted (p < 0.05). Data from *IL2* validation screen, n=3 biological replicates, 8 gRNAs per enhancer. P values are calculated by Wilcoxon test unless noted otherwise. Significance cutoffs: ns p > 0.05, * p <= 0.05, ** p <= 0.01, *** p <= 0.001, **** p <= 0.0001.

**Supplementary Fig. S10. (related to Fig. 4).**
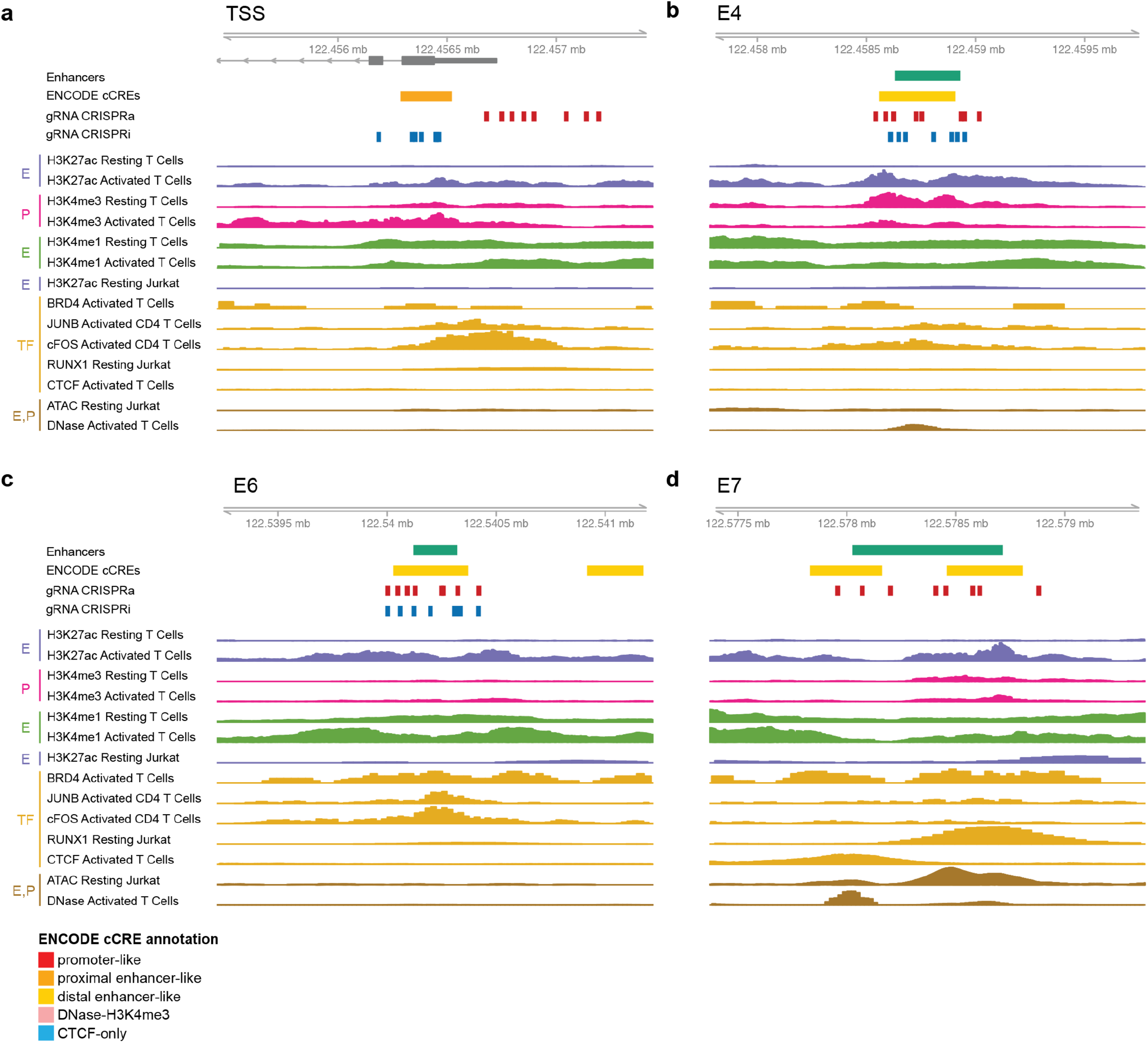
Genome tracks for strong functional enhancers of *IL2*. Genome tracks for selected regions of *IL2* locus: (a) TSS, (b) E4, (c) E6, and (d) E7. Same tracks as shown in **Fig. 3a** and **Supplementary Fig. S8a**, with additional track: ENCODE candidate cis-regulatory elements combined from all cell types (cCREs)^64,65^.

## METHODS

### Cell Culture

Lenti-X HEK293T (Clontech) cells were cultured in DMEM (Gibco) with L-glutamine and sodium pyruvate supplemented with 10% FBS (Gibco) and 1% penicillin-streptomycin (pen-strep, Gibco), and passaged using TrypLE Express (Gibco). K562 (ATCC) cultured in RPMI 1640 (Gibco) with L-glutamine supplemented with 10% FBS and 1% pen-strep. Jurkat Clone E6-1 cells (ATCC) were cultured in RPMI 1640 with L-glutamine (Gibco) supplemented with 10% FBS, 10mM HEPES (Gibco), 1mM sodium pyruvate (Gibco), and 1% pen-strep. Cells were routinely tested for mycoplasma using MycoAlert PLUS Detection Kit (Lonza) and found to be negative.

### CRISPRai Construct Generation

The CRISPRai construct was cloned in the format: TRE3G-VPR-dSaCas9-P2A-dSpCas9-BFP-KRAB-EF1a-Bleo-T2A-rtTA). The vector containing the TRE3G and TetOn system was PiggyBac; the zeocin resistance gene and the TetOn 3G transactivator were driven by the EF1a promoter (gifted by Stanley Qi lab)^86^. The Super PiggyBac Transposase plasmid was obtained from Systems Biosciences. VPR was obtained from pSLQ2349 (gifted by Stanley Qi lab), dSaCas9 was obtained from pSLQ2840 (Addgene 84246), dSpCas9-BFP-KRAB was obtained from pHR-SFFV-dCas9-BFP-KRAB (Addgene 46911). Constructs were cloned using Gibson Assembly (NEBuilder HIFI DNA Assembly) and confirmed by sanger sequencing (Elim Biopharm). Primers and oligos were obtained from Elim Biopharm and IDT DNA Technologies.

### CRISPR gRNA Cloning

Primers and oligos for bulk validation experiments were obtained from Elim Biopharm and IDT. All cloned plasmids were confirmed by sanger sequencing (Elim Biopharm). Individual single gRNAs were cloned using Gibson Assembly (NEBuilder HIFI DNA Assembly). For validation and Perturb-seq experiments, gRNAs were constructed from pSLQ2853-3 pHR: U6-Sasgv2CXCR4-1 CMV-EGFP (Addgene 84254) and pSLQ1852-2 pHR: U6-SpsgCD95-1 CMV-EGFP (Addgene 84151) with slight modifications. For dSaCas9 gRNAs, GFP was replaced with mScarlet (pmScarlet_Giantin_C1, Addgene 85048).

For Perturb-seq single gRNAs, gRNAs pools were constructed from 2 gRNA backbones, one with the dSpCas9 gRNA scaffold and the other with dSaCas9 gRNA scaffold. Pools were cloned in an arrayed format by ordering top and bottom ∼31-33bp gRNA oligos from IDT with appropriate overhangs. Top and bottom oligos were combined at a final concentration of 100mM in annealing buffer composed of 100mM potassium acetate, 30mM HEPES-KOH pH 7.4, and 2mM magnesium acetate in water (adapted from Jonathan Weismann lab protocols, https://weissman.wi.mit.edu/resources/) annealed on a thermocycler at 95C 4min, then cooled slowly in insulation for 3 hours. Annealed oligos were pooled, phosphorylated using T4 PNK (NEB) at 37C 30min with 65C 20min PNK inactivation, then ligated into the previously digested and dephosphorylated (Fast AP, Thermo Fisher Scientific) lentiviral gRNA backbone using T4 ligase (NEB). Pools were transformed by heat shock into Stbl3 competent cells (Thermo Fisher Scientific).

For Perturb-seq double gRNAs, gRNA pools were constructed in a two-step cloning process with the final gRNA cassette format: [mu6-gRNA1-gRNA scaffold1-hu6-gRNA2-gRNA scaffold2] (**Supplementary Fig. S2a**). Oligo pools (IDT) containing ∼200bp oligos with the format amplification primer-digest site-gRNA1-scaffold1-hu6 landing pad-digest site-amplification primer. For step 1, oligo pools were PCR amplified in multiple reactions with low cycle number (NEB Ultra II MM), digested, and size selected via gel purification (EGel EX, Thermo Fisher Scientific). Inserts were ligated into predigested gRNA backbones with T4 ligase overnight 16C 16hrs and inactivated 65C 10min, transformed into Stbl3 competent cells and grown at 30C. For step 2, plasmid products were digested, dephosphorylated, and gel size selected, and the previously digested hu6 PCR fragment (from pMJ117 Addgene 85997) with appropriate overhangs was inserted via T4 ligation. Original vector backbone and intermediate backbone product were designed for digestion with Esp3I (BsmBI, NEB), inserts were designed for digestion with BsaI (NEB).

For the enhancer gRNA enrichment screen, double gRNA pools were constructed in a one-step cloning process and the final format was [mu6-gRNA1-Sp gRNA scaffold-hu6-gRNA2-Sa gRNA scaffold] (**Supplementary Fig. S2b**). Primer pools were obtained from IDT and contained gRNA sequences as well as priming sequences complementary to the dSpCas9 gRNA scaffold and the hu6 promoter.

Primers were used to generate a PCR product in the format of [mu6 fragment-gRNA1-Sp gRNA scaffold-hu6-gRNA2-Sa gRNA scaffold fragment], flanked by BsmBI digestion sites. The PCR product and backbone were digested separately and ligated with T4 ligase following recommended protocols.

### Stable Cell Line Generation

Stable cell lines were generated by electroporation via the Neon Transfection System (Thermo Fisher Scientific). Cells were electroporated and allowed to recover in fresh media for 3 days, selected with zeocin (Thermo Fisher Scientific) for 10 days, then analyzed by flow cytometry for BFP to confirm dCas9 cassette expression near 100% of cells. Electroporation parameters recommended by the Neon system were used for the respective cell types.

### qPCR

Primers were obtained from Elim Biopharm. Brilliant II SYBR Green qPCR MM (Agilent Technologies) and was used. Primers were validated prior to use by examining the melt curve. Analysis was performed using the double delta Ct method, relative to the housekeeping gene *ACTB* and NTC gRNA controls.

### Lentivirus Production

Lenti-X HEK293T cells were seeded on plates the night before, and then transfected with packaging plasmids psPAX2 (1.5ug, Addgene 12260) and pMD2.G (4.5ug, Addgene 12259), and viral expression vector (6ug) per 10cm plate using Opti-MEM (Gibco) and Lipofectamine 3000 reagents (Thermo Fisher Scientific). Viral supernatant was collected at 48 hours and concentrated 100x using Lenti-X Concentrator (Clontech), resuspended in cell culture media, and stored at −80C.

### Flow Cytometry and FACS for Cell-Surface Markers and Intracellular Cytokines

For validation CRISPRi experiments, cells were stained with CD3E-BV785 (Biolegend clone OKT3) or CD47-BV605 antibodies (Biolegend clone CC2C6) for 30min at 4C. For sorting, all cells for Perturb-seq gRNA enrichment screens were stained with Zombie NIR fixable viability dye 1:1000 dilution in PBS (Biolegend). Cells were analyzed by flow cytometry (Attune Nxt, Thermo Fisher Scientific, or LSR II, BD Biosciences), or sorted based on stained markers and gRNA expression (GFP or mScarlet) (FACS Aria II, BD Biosciences). All cells were stained in flow cytometry staining buffer (eBioscience). Fluorescence-activated cell sorting (FACS) was performed at the Stanford Shared FACS Facility.

For Jurkat intracellular cytokine staining, activated cells were stained with Zombie NIR viability dye at 1:1000 dilution in PBS at 10M cells / 100ul for 15min at 4C, washed with staining buffer, and fixed using Cyto-Fast Fix Perm Buffer Set (Biolegend) for 25min at 22C, washed with cell staining buffer, and stored in Cyto-Last Buffer (Biolegend) at 4C in the dark for 1-3 days. Before sorting, fixed cells were permeabilized and stained with IL2-BV711 (Biolegend clone MQ1-17H12) and IFNG-APC (Biolegend clone B27) antibodies for 45min at 22C, washed with fix/perm buffer, and resuspended in staining buffer.

### Pooled K562 Single-Cell Screening

Cells were infected with lentivirus gRNA pools at an MOI of 0.1, as confirmed by flow cytometry for GFP or mScarlet expression on days 2 and 3 post-infection. Polybrene was added at the time of infection at 8ug/ml and cells were changed to fresh media the next day. Doxycycline (Dox) was added at the time of infection and refreshed every 24 hours. Cells were expanded for 6 days post-infection, and frozen in aliquots on day 6 in Cryotstor CS10 (StemCell Technologies). Before sorting, cells were thawed and allowed to recover in culture in dox+ media for 18 hrs. Cells were sorted by FACS for live, gRNA+ cells.

### Single-Cell Library Preparation

Sorted cells were prepared using the Chromium Next GEM Single Cell 5’ Kit v2, Chromium Next GEM Chip K Single Cell Kit, and Library Construction Kit (10x Genomics), following the Chromium Next GEM Single Cell 5’ Reagent Kits v2 (Dual Index) with Feature Barcoding user guide (CG000330 Rev A).

GEX libraries were constructed as recommended. Modifications to the protocol for gRNA library construction were as follows: oligos complementary to each of the gRNA scaffolds (Sa and Sp) were spiked into the reverse-transcription (RT) reaction at 11.43 pmol each.

Sa: AAGCAGTGGTATCAACGCAGAGTACacaagttgacgagataaacacgg Sp: AAGCAGTGGTATCAACGCAGAGTACcgactcggtgccactttttc

For step 2.2, cDNA primers were used (green, 10x PN 2000089) instead of feature cDNA primers (purple, 10x PN 2000277). For step 2.3, GEX is in the pellet (2.3A) and gRNAs are in the supernatant (2.3B), both portions were retained and library construction was performed separately. gRNA library construction was performed using a custom PCR protocol, and Sa and Sp gRNA libraries were constructed separately. PCR1: outer nested PCR, F CTACACGACGCTCTTCCGATCT, R_sa acaagttgacgagataaacacgg, R_sp CGACTCGGTGCCACTTTTTC (98C 3min; 20 cycles 98C 20s, 66C(Sa)/68C(Sp) 30s, 72C 20s; 72C 5min). PCR2: inner nested PCR and adapter common region addition, F same primer as PCR1, R_sa GTGACTGGAGTTCAGACGTGTGCTCTTCCGATCTgataaacacggcattttgccttg, R_sp GTGACTGGAGTTCAGACGTGTGCTCTTCCGATCTcaagttgataacggactagcctt (same cycling conditions as PCR1, with annealing temperatures 66C(Sa)/65C(Sp)). PCR3: sample index PCR, P5 and P7 Dual Index TT Set A (98C 3min; 15 cycles 98C 20s, 54C 30s, 72C 1min; 72C 5min). After each PCR, products were run on EGel EX 2% agarose and size selected.

### Pooled Jurkat Screening

Cells were infected on day 0 with lentivirus gRNA pools at an MOI of 0.2, as confirmed by flow cytometry for mScarlet expression on days 2 and 3 post-infection. Cells were infected in 8ug/ml polybrene overnight, and next morning changed to fresh media. On day 3, 0.5ug/ml puromycin (puro, Thermo Fisher Scientific) was added, cells were selected for gRNA expression for 4 days, and confirmed by flow cytometry to have near 100% gRNA expression post-selection. On day 7, puro selection was stopped and dox induction (final concentration 1ug/ml) of the CRISPRai dCas9 construct was started. Dox was refreshed every 24 hrs, for a total of 6 days of dox induction. On day 13, cells were activated at ∼2-4M cells/ml for 8 hrs using CD3 antibody (Biolegend clone OKT3) coated tissue culture plates, and media containing dox (1ug/ml), CD28 antibody (3ug/ml, Biolegend clone CD28.2),

PMA (1x), ionomycin (1x), Brefeldin A (1x) (PMA/iono/BrefA were used from Cell Activation Cocktail, Biolegend). Culture plates were coated with CD3 antibody diluted in PBS at 5ug/ml for 1.5hrs at 37C, then rinsed gently with media before adding cells. gRNA+ cell number (accounting for MOI) was maintained at 1000x the number of gRNAs included in the gRNA pool throughout the screen. For the validation screen, the above steps were followed exactly. For the primary screen, the above steps were modified to begin dox-induction at the time infection, puro selection was performed from day 3-7, and cells were activated and the screen was stopped on day 7.

### CRISPR gRNA Enrichment Library Preparation

Genomic DNA (gDNA) was extracted from sorted cells for different cytokine populations. Primary screen: IL2+IFNG-(IL2), IFNG+IL2-(IFNG), IL2+IFNG+ (DP, double positive), IL2-IFNG-(NEG), and unsorted cells (UN). Validation screen: IL2+ (IL2), IL2-(NEG), and unsorted cells (UN). Cells were washed with PBS, and resuspended in 1x lysis buffer (10mM Tris pH 8, 5mM EDTA, 0.5% SDS, 1x (0.4mg/ml) Proteinase K (Thermo Fisher Scientific) in water at 10M cells/800ul, incubated at 55C 2hrs, then 65C 16-20hrs overnight. Samples were then cooled to room temperature for 10min and Triton X-100 (Sigma Aldrich) was added to a final concentration of 0.5%. The number of cells per population used for gDNA extraction was 0.2-15M for the primary screen and 10-20M for the validation screen. For samples >2M sorted cells, gDNA was then purified using the Quick-DNA Miniprep kit (Zymo), following the “Cell Suspensions and Proteinase K Digested Samples” recommended protocol. For samples <2M cells, a precipitate-based method was used for gDNA extraction; briefly, following addition of Triton X-100, sodium acetate was added to a final concentration of 10%, 2.5x volumes of 100% EtOH was added, samples were placed at −20C for 1hr followed by centrifuge at 20,000xg 15min at 4C, supernatant was removed, 75% EtOH was added to wash, centrifuged again, pellets were dried overnight at room temperature, and resuspended in elution buffer.

Library preparation from gDNA was performed by 3 PCR steps. PCR1: multiple reactions per sample were set up with 2ug or less of gDNA with outer nested primers complementary to the gRNA cassette (98C 3min; 14 cycles of 98C 20s, 58C 20s, 72C 40s; 72C 2min) and concentrated with DNA Clean and Concentrator (Zymo). PCR2: inner nested primers (98C 30s; 6 cycles of 98C 15s, 60C 15s, 72C 45s; 72C 2min) and size selected using SPRI beads 0.75x cleanup. PCR3: Tru-seq based indexing primers (98C 30s; 6 cycles of 98C 15s, 63C 15s, 72C 45s; 72C 2min) and size selected using SPRI beads 0.75x cleanup. After each PCR, products were checked on EGel EX 2% agarose.

Primer sequences:

PCR1

mU6_outer_fw cagcacaaaaggaaactcaccctaactgtaaag

sasgRNA_PCR_3Rev tctcgccaacaagttgacgagataaaca

PCR2

p7_saRNA_stagger2_rev GTGACTGGAGTTCAGACGTGTGCTCTTCCGATCTccttgttatagtagattctgtttccagagtactaTAAC

p7_saRNA_stagger1_rev GTGACTGGAGTTCAGACGTGTGCTCTTCCGATCTcttgttatagtagattctgtttccagagtactaTAAC

p7_saRNA_stagger0_rev GTGACTGGAGTTCAGACGTGTGCTCTTCCGATCTtgttatagtagattctgtttccagagtactaTAAC

p5_mU6_0nt_stagger ACACTCTTTCCCTACACGACGCTCTTCCGATCTtcccttggagaaccaccttgt

p5_mU6_1nt_stagger ACACTCTTTCCCTACACGACGCTCTTCCGATCTCtcccttggagaaccaccttgt

p5_mU6_2nt_stagger ACACTCTTTCCCTACACGACGCTCTTCCGATCTGCtcccttggagaaccaccttgt

### Sequencing

Library quality was checked by Bioanalyzer (Agilent) and quantified by KAPA Library Quantification Kit (Roche). Sequencing was performed on NovaSeq 6000 (Illumina, Novogene) or NextSeq 550 (Illumina). For single-cell Perturb-seq libraries, libraries were sequenced at ∼6,000 reads/cell for gRNA and ∼30,000-50,000 reads/cell for GEX. For the Jurkat enhancer screens, gRNA enrichment libraries were sequenced at ∼1.5M reads per sample for the primary screen and ∼7.5M reads per sample for the validation screen (∼1200 reads/gRNA post-filtering), and gRNA1 and gRNA2 in the double gRNA cassette were sequenced in R1 and R2 paired reads, respectively, and then paired in silico.

### Single-Cell gRNA and Transcriptome Analysis

The scRNA-seq reads were aligned to GRCh38 genome and quantified using cellranger count (10x Genomics). gRNA reads were processed using cellranger count with gRNA sequences as the feature reference. Later analyses were all performed in R using Seurat.

Data from 5 total captures was combined to one Seurat object. Cells were filtered: number of genes > 200, number of genes < 5500-8300, transcriptome UMIs < 27000-75000, percent mitochondrial reads < 10%, detected gRNAs > 20 (background signal distribution), gRNA UMIs > 50, with exact parameters differing for each capture. gRNA labels for each cell were assigned based on cellranger feature calls.

Only cells with 1 or 2 cellranger-detected gRNAs were retained for single gRNA and double gRNA captures, respectively. gRNA groups with < 250 cells with target gene detected or with low cell numbers (n < 20) were removed. gRNA pools contained 2 gRNAs per gene for single perturbations and 1 gRNA per gene for double perturbations. For each gene, the gRNA with higher magnitude log2FC in the single perturbations was used for the double perturbation gRNA. If the 2 gRNAs for a given gene were not concordant in target gene log2FC expression for single perturbations, only the gRNA with greater magnitude change was retained for analysis. For these reasons, the following gRNAs were removed from the dataset: CEBPA-a1, CEBPA-a2, MAP2K3-a1, MYC-a1, MYC-a2, MYC-e1-a1, MYC-e1-a2, SPI1-a2, RIPK2-a2, ATF5-i1, CEBPB-i1, FOSL1-i2. Gene expression was log normalized with a scaling factor of 1e4. gRNA expression was normalized using relative counts with a scaling factor of 100.

For Fig. 1, only gRNA groups with > 40 cells were included. Differential expression for CRISPR target genes was performed FindMarkers() using normalized counts and a logistic regression model with batch as a latent variable. Batch was defined as the day on which 10x captures were performed, either day 1 or day 2. For Fig. 2, the top 2000 most variable genes were found using variance stabilization transformation (vst). All genes were centered and scaled, and batch and percent mitochondrial reads were regressed out using ScaleData(). PCA was performed on the top 2000 most variable genes, followed by nearest neighbor graph construction, cluster determination using the original Louvain algorithm, and UMAP dimensionality reduction using the top PCs. All further analyses were performed with regression on batch as the only latent variable except for UMAP reduction of 24,661 cells in Supplementary Fig. S3a, which was regressed on batch and percent mitochondrial reads.

Next, the subset of cells in the SPI1 and GATA1 gRNA groups were retained, and variable gene selection was repeated. Perturbation-driven cells were identified as clusters that were not composed of equal representations from all gRNA groups. Non-perturbation driven cells were removed and variable feature selection, PCA, neighbor graph construction, clustering, and UMAP reduction were performed again. All differential expression testing was performed on either all genes or genes in the indicated TF target gene sets using normalized counts and logistic regression with batch as a latent variable. For module score analysis, ENCODE TF target gene sets for SPI1 and GATA1 were downloaded from Harmonizome^64-66^ and genes were identified as being unique to either set or shared. Module scores were calculated using AddModuleScore() using normalized, scaled, and batch regressed counts and statistical testing was performed using a Wilcoxon test.

All functions referenced above are from Seurat. Statistical testing was performed using stat_compare_means() from ggpubr or FindMarkers() and FindAllMarkers() from Seurat. All plots were generated in R using Seurat, ggplot, ggpubr, pheatmap.

### CRISPR gRNA Enrichment Analysis

gRNA reads were processed in Python to perform in silico pairing of gRNA1 and gRNA2 from paired end reads and to generate a raw read counts per gRNA matrix. All later analyses were performed in R. gRNA pairs were filtered for pairs with the sum of raw read counts across all sorted populations > 300 reads. Reads were normalized per sample by dividing by the total reads per sample and scaling by 1e6, and log2 transformed with a pseudocount of 1. Fold change was calculated between each population versus the cytokine-negative population (NEG). Z-scores were computed by centering and scaling relative to the mean and standard deviation of all NTC gRNAs. Z-scores were used for all further analyses. Z-scores were calculated independently for the primary and validation *IL2* locus screens.

Expected double gRNA enrichment was calculated by summing the log2FC gRNA enrichment z-scores of the corresponding singles: log2FC z-score(single1) + log2FC z-score(single2). Residuals were calculated from the line of best fit between expected and observed double log2FC z-score. The Fisher’s Exact test was performed with fisher.test() by counting the occurrences of gRNAs for a given enhancer in the top and bottom 5% of residuals for the expected vs observed double log2FC gRNA enrichment fit. Percent of TSS perturbation for enhancer gRNAs was calculated for CRISPRa using true fold change gRNA enrichment (IL2+ / IL2-) in single perturbations by (FC E_a_gRNA) / (FC TSS_a_gRNA), and for CRISPRi by (1 - FC E_i_gRNA) / (1 - FC TSS_i_gRNA).

For histone ChIP-Seq analysis, bigWig files containing “fold change over control” were downloaded from ENCODE^64,65^. File accessions used were: for activated T cells ENCFF233LPC ENCFF370YXG ENCFF356ZKI ENCFF704NYS ENCFF741XLV ENCFF158HYB ENCFF232FZK ENCFF206YVE ENCFF336KWY ENCFF164WIU ENCFF060VND ENCFF398QTX ENCFF940OQY ENCFF903VVJ ENCFF356TWG ENCFF248VJB ENCFF690AHR ENCFF243FBP ENCFF624BMC ENCFF352EYP and for resting T cells ENCFF906URN ENCFF787PDH ENCFF787LLC ENCFF820GJE ENCFF984ZEE ENCFF829WQD ENCFF055UPO ENCFF459VQV ENCFF041OBG ENCFF543OQM ENCFF863YFO ENCFF896VDJ ENCFF560YNU ENCFF309ISK ENCFF953MIX ENCFF478JER. Peaks overlapping each enhancer were used to estimate enhancer-specific histone signatures using GRanges and IRanges. For TF motif enrichment analysis, PFMs were downloaded from JASPAR^79^: JASPAR2022_CORE_vertebrates_non-redundant_pfms_jaspar.txt. TF motif enrichment in each enhancer was performed using matchMotifs() from motifmatchr^87^ using parameters: genome=hg38, out=scores, bg=subject, p.cutoff=5e-5 and filtered for the top scoring motifs.

For genome tracks, the following datasets were used. Gene region: TxDb.Hsapiens.UCSC.hg38.knownGene. ABC model predictions: AllPredictions.AvgHiC.ABC0.015.minus150.ForABCPaperV3.txt.gz^42^. The following file accessions were downloaded from ENCODE: H3K27ac activated T cell ChIP-seq ENCFF356ZKI, H3K27ac resting T cell ChIP-seq ENCFF820GJE, H3K4me3 activated T cell ChIP-seq ENCFF940OQY, H3K4me3 resting T cell ChIP-seq ENCFF863YFO, H3K4me1 activated T cell ChIP-seq ENCFF755MCS, H3K4me1 resting T cell ChIP-seq ENCFF041OBG, activated T cells DNase-seq ENCFF997BFO, CTCF activated T cell ChIP-seq ENCFF523IEI^64,65^. H3K27ac resting Jurkat ChIP-seq (GEO GSM1697882)^38^; BRD4 activated T cell ChIP-seq GSM5573170_Stim_BRD4.bw (GEO GSM5573170)^84^; JUNB and cFOS activated CD4 T cell ChIP-seq (GEO GSE116695, SRA SRR7475866 and SRR7475865)^80^, RUNX1 resting Jurkat ChIP-seq (GEO GSM1697879)^38^; resting Jurkat ATAC-seq (GEO GSM4130892)^85^. Fastq files downloaded from SRA were converted to BigWig files using Galaxy tools and recommended pipelines^88^. The ENCODE cCRE track was generated using Gviz function UcscTrack(track = “encodeCcreCombined”.).

For SRE score analysis, enhancer coordinates and SRE scores were downloaded from the Multiplexed CRISPRi EnhancerNet website^36^ for the *IL2* gene in Jurkat T cells. For the subset of enhancers shared between our screen and the SRE dataset, SRE score was plotted for all enhancer pairs. The following enhancers were shared between the CRISPRai screen and the SRE dataset: E4, E5, E7, E8, E9.

All plots were generated in R using ggplot, ggpubr, pheatmap. Genome tracks were generated using rtracklayer and Gviz. In all boxplots, statistical analysis was performed using stat_compare_means() from ggpubr. Statistical significance in log2FC gRNA enrichment was performed using a two-sided Wilcoxon test using wilcox.test() and corrected for multiple testing using the Benjamini-Hochberg procedure.

## ACKNOWLEDGEMENTS

We thank Stanley Qi lab for gift of plasmids, Profs. A. Marson, J. Engreitz and S. Qi for discussion. We also thank J. A. Belk, S. K. Wang, R. Chen, K. E. Yost, K. Kraft, C. K. Chen for helpful discussions and technical advice. Supported by the National Science Foundation Graduate Research Fellowship (N.M.P), NIH RM1-HG007735 and Scleroderma Research Foundation (to H.Y.C.). H.Y.C. is an Investigator of the Howard Hughes Medical Institute.

## AUTHOR CONTRIBUTIONS

N.M.P. and H.Y.C. conceived the project. N.M.P performed experiments and computational analysis for CRISPRai method development, method validation and optimization, K562 Perturb-seq screen, and Jurkat gene regulation screens. Q.S. cloned gRNA pools for Jurkat screens and advised on computational analysis. K.R.P. was involved in early discussions of project design. N.M.P. and H.Y.C. wrote the manuscript with input from all authors.

## SUPPLEMENTARY TABLES

**Supplementary Table S1. Single gRNA sequences for K562 screen**.

**Supplementary Table S2. Double gRNA sequences for K562 screen**.

**Supplementary Table S3. gRNA sequences for Jurkat *IL2* primary screen**.

**Supplementary Table S4. gRNA sequences for Jurkat *IFNG* primary screen**.

**Supplementary Table S5. gRNA sequences for Jurkat *IL2* validation screen**.

## REFERENCES

1. Maeder, M. L. et al. CRISPR RNA-guided activation of endogenous human genes. Nat Methods 10, 977–979 (2013).

2. Perez-Pinera, P. et al. RNA-guided gene activation by CRISPR-Cas9-based transcription factors. Nat Methods 10, 973–976 (2013).

3. Tanenbaum, M. E., Gilbert, L. A., Qi, L. S., Weissman, J. S. & Vale, R. D. A protein-tagging system for signal amplification in gene expression and fluorescence imaging. Cell 159, 635–646 (2014).

4. Chavez, A. et al. Highly efficient Cas9-mediated transcriptional programming. Nat Methods 12, 326–328 (2015).

5. Konermann, S. et al. Genome-scale transcriptional activation by an engineered CRISPR-Cas9 complex. Nature 517, 583–588 (2015).

6. Qi, L. S. et al. Repurposing CRISPR as an RNA-Guided Platform for Sequence-Specific Control of Gene Expression. Cell 152, 1173–1183 (2013).

7. Gilbert, L. A. et al. CRISPR-Mediated Modular RNA-Guided Regulation of Transcription in Eukaryotes. Cell 154, 442–451 (2013).

8. Gilbert, L. A. et al. Genome-Scale CRISPR-Mediated Control of Gene Repression and Activation. Cell 159, 647–661 (2014).

9. Liu, S. J. et al. CRISPRi-based genome-scale identification of functional long noncoding RNA loci in human cells. Science 355, eaah7111 (2017).

10. Joung, J. et al. Genome-scale activation screen identifies a lncRNA locus regulating a gene neighbourhood. Nature 548, 343–346 (2017).

11. Kampmann, M. CRISPRi and CRISPRa Screens in Mammalian Cells for Precision Biology and Medicine. ACS Chem. Biol. 13, 406–416 (2018).

12. Klann, T. S. et al. CRISPR-Cas9 epigenome editing enables high-throughput screening for functional regulatory elements in the human genome. Nat Biotechnol 35, 561–568 (2017).

13. Simeonov, D. R. et al. Discovery of stimulation-responsive immune enhancers with CRISPR activation. Nature 549, 111–115 (2017).

14. Fulco, C. P. et al. Systematic mapping of functional enhancer-promoter connections with CRISPR interference. Science 354, 769–773 (2016).

15. Schraivogel, D. et al. Targeted Perturb-seq enables genome-scale genetic screens in single cells. Nat Methods 17, 629–635 (2020).

16. Du, D. et al. Genetic interaction mapping in mammalian cells using CRISPR interference. Nat Methods 14, 577–580 (2017).

17. Han, K. et al. Synergistic drug combinations for cancer identified in a CRISPR screen for pairwise genetic interactions. Nat Biotechnol 35, 463–474 (2017).

18. Boettcher, M. et al. Dual gene activation and knockout screen reveals directional dependencies in genetic networks. Nat Biotechnol 36, 170–178 (2018).

19. Gasperini, M. et al. A Genome-wide Framework for Mapping Gene Regulation via Cellular Genetic Screens. Cell 176, 377-390.e19 (2019).

20. Norman, T. M. et al. Exploring genetic interaction manifolds constructed from rich single-cell phenotypes. Science 365, 786–793 (2019).

21. Joung, J. et al. Genome-scale CRISPR-Cas9 knockout and transcriptional activation screening. Nature Protocols 12, 828–863 (2017).

22. Dahlman, J. E. et al. Orthogonal gene knockout and activation with a catalytically active Cas9 nuclease. Nat Biotechnol 33, 1159–1161 (2015).

23. Najm, F. J. et al. Orthologous CRISPR-Cas9 enzymes for combinatorial genetic screens. Nat Biotechnol 36, 179–189 (2018).

24. Lin, Y. et al. CRISPR/Cas9 systems have off-target activity with insertions or deletions between target DNA and guide RNA sequences. Nucleic Acids Res 42, 7473–7485 (2014).

25. Tsai, S. Q. et al. GUIDE-seq enables genome-wide profiling of off-target cleavage by CRISPR-Cas nucleases. Nat Biotechnol 33, 187–197 (2015).

26. Aguirre, A. J. et al. Genomic Copy Number Dictates a Gene-Independent Cell Response to CRISPR/Cas9 Targeting. Cancer Discov 6, 914–929 (2016).

27. Morgens, D. W. et al. Genome-scale measurement of off-target activity using Cas9 toxicity in high-throughput screens. Nat Commun 8, 15178 (2017).

28. Shin, H. Y. et al. CRISPR/Cas9 targeting events cause complex deletions and insertions at 17 sites in the mouse genome. Nat Commun 8, 15464 (2017).

29. Kosicki, M., Tomberg, K. & Bradley, A. Repair of double-strand breaks induced by CRISPR-Cas9 leads to large deletions and complex rearrangements. Nat Biotechnol 36, 765–771 (2018).

30. Lian, J., HamediRad, M., Hu, S. & Zhao, H. Combinatorial metabolic engineering using an orthogonal tri-functional CRISPR system. Nat Commun 8, 1688 (2017).

31. Martella, A. et al. Systematic Evaluation of CRISPRa and CRISPRi Modalities Enables Development of a Multiplexed, Orthogonal Gene Activation and Repression System. ACS Synthetic Biology (2019) doi:10.1021/acssynbio.8b00527.

32. Black, J. B. et al. Master Regulators and Cofactors of Human Neuronal Cell Fate Specification Identified by CRISPR Gene Activation Screens. Cell Reports 33, 108460 (2020).

33. Wu, F., Shim, J., Gong, T. & Tan, C. Orthogonal tuning of gene expression noise using CRISPR-Cas. Nucleic Acids Research 48, e76 (2020).

34. Jensen, T. I. et al. Targeted regulation of transcription in primary cells using CRISPRa and CRISPRi. Genome Res gr.275607.121 (2021) doi:10.1101/gr.275607.121.

35. Ameruoso, A., Villegas Kcam, M. C., Cohen, K. P. & Chappell, J. Activating natural product synthesis using CRISPR interference and activation systems in Streptomyces. Nucleic Acids Research 50, 7751–7760 (2022).

36. Lin, X. et al. Nested epistasis enhancer networks for robust genome regulation. Science 377, 1077–1085 (2022).

37. Hnisz, D. et al. Convergence of developmental and oncogenic signaling pathways at transcriptional super-enhancers. Mol Cell 58, 362–370 (2015).

38. Hnisz, D. et al. Activation of proto-oncogenes by disruption of chromosome neighborhoods. Science 351, 1454–1458 (2016).

39. Catarino, R. R. & Stark, A. Assessing sufficiency and necessity of enhancer activities for gene expression and the mechanisms of transcription activation. Genes Dev 32, 202–223 (2018).

40. Haberle, V. & Stark, A. Eukaryotic core promoters and the functional basis of transcription initiation. Nat Rev Mol Cell Biol 19, 621–637 (2018).

41. Fulco, C. P. et al. Activity-by-contact model of enhancer-promoter regulation from thousands of CRISPR perturbations. Nature Genetics 51, 1664–1669 (2019).

42. Nasser, J. et al. Genome-wide enhancer maps link risk variants to disease genes. Nature 593, 238–243 (2021).

43. Neumayr, C. et al. Differential cofactor dependencies define distinct types of human enhancers. Nature 606, 406–413 (2022).

44. Zuin, J. et al. Nonlinear control of transcription through enhancer-promoter interactions. Nature 604, 571–577 (2022).

45. Horlbeck, M. A. et al. Compact and highly active next-generation libraries for CRISPR-mediated gene repression and activation. eLife 5, e19760 (2016).

46. Tian, R. et al. Genome-wide CRISPRi/a screens in human neurons link lysosomal failure to ferroptosis. Nat Neurosci 1–15 (2021) doi:10.1038/s41593-021-00862-0.

47. Schmidt, R. et al. CRISPR activation and interference screens decode stimulation responses in primary human T cells. Science (2022) doi:10.1126/science.abj4008.

48. Adamson, B. et al. A Multiplexed Single-Cell CRISPR Screening Platform Enables Systematic Dissection of the Unfolded Protein Response. Cell 167, 1867-1882.e21 (2016).

49. Dixit, A. et al. Perturb-Seq: Dissecting Molecular Circuits with Scalable Single-Cell RNA Profiling of Pooled Genetic Screens. Cell 167, 1853-1866.e17 (2016).

50. Jaitin, D. A. et al. Dissecting Immune Circuits by Linking CRISPR-Pooled Screens with Single-Cell RNA-Seq. Cell 167, 1883-1896.e15 (2016).

51. Datlinger, P. et al. Pooled CRISPR screening with single-cell transcriptome read-out. Nat Methods 14, 297–301 (2017).

52. Xie, S., Duan, J., Li, B., Zhou, P. & Hon, G. C. Multiplexed Engineering and Analysis of Combinatorial Enhancer Activity in Single Cells. Molecular Cell 66, 285-299.e5 (2017).

53. Rekhtman, N., Radparvar, F., Evans, T. & Skoultchi, A. I. Direct interaction of hematopoietic transcription factors PU.1 and GATA-1: functional antagonism in erythroid cells. Genes Dev. 13, 1398–1411 (1999).

54. Zhang, P. et al. PU.1 inhibits GATA-1 function and erythroid differentiation by blocking GATA-1 DNA binding. Blood 96, 2641–2648 (2000).

55. Graf, T. & Enver, T. Forcing cells to change lineages. Nature 462, 587–594 (2009).

56. Burda, P. et al. GATA-1 Inhibits PU.1 Gene via DNA and Histone H3K9 Methylation of Its Distal Enhancer in Erythroleukemia. PLOS ONE 11, e0152234 (2016).

57. Nishimasu, H. et al. Crystal Structure of Staphylococcus aureus Cas9. Cell 162, 1113–1126 (2015).

58. Friedland, A. E. et al. Characterization of Staphylococcus aureus Cas9: a smaller Cas9 for all-in-one adeno-associated virus delivery and paired nickase applications. Genome Biology 16, 257 (2015).

59. Mimitou, E. P. et al. Multiplexed detection of proteins, transcriptomes, clonotypes and CRISPR perturbations in single cells. Nat Methods 16, 409–412 (2019).

60. Replogle, J. M. et al. Combinatorial single-cell CRISPR screens by direct guide RNA capture and targeted sequencing. Nat Biotechnol 38, 954–961 (2020).

61. Qian, Y., Huang, H.-H., Jiménez, J. I. & Del Vecchio, D. Resource Competition Shapes the Response of Genetic Circuits. ACS Synth. Biol. 6, 1263–1272 (2017).

62. Chen, P.-Y., Qian, Y. & Del Vecchio, D. A Model for Resource Competition in CRISPR-Mediated Gene Repression. in 2018 IEEE Conference on Decision and Control (CDC) 4333–4338 (2018). doi:10.1109/CDC.2018.8619016.

63. McCarty, N. S., Graham, A. E., Studená, L. & Ledesma-Amaro, R. Multiplexed CRISPR technologies for gene editing and transcriptional regulation. Nat Commun 11, 1281 (2020).

64. ENCODE Project Consortium. The ENCODE (ENCyclopedia Of DNA Elements) Project. Science 306, 636–640 (2004).

65. ENCODE Project Consortium. A user’s guide to the encyclopedia of DNA elements (ENCODE). PLoS Biol 9, e1001046 (2011).

66. Rouillard, A. D. et al. The harmonizome: a collection of processed datasets gathered to serve and mine knowledge about genes and proteins. Database 2016, baw100 (2016).

67. Subramanian, A. et al. Gene set enrichment analysis: A knowledge-based approach for interpreting genome-wide expression profiles. Proceedings of the National Academy of Sciences 102, 15545–15550 (2005).

68. Liberzon, A. et al. Molecular signatures database (MSigDB) 3.0. Bioinformatics 27, 1739–1740 (2011).

69. Rubin, A. J. et al. Coupled Single-Cell CRISPR Screening and Epigenomic Profiling Reveals Causal Gene Regulatory Networks. Cell 176, 361-376.e17 (2019).

70. Warner, J. L. & Arnason, J. E. Alemtuzumab use in relapsed and refractory chronic lymphocytic leukemia: a history and discussion of future rational use. Ther Adv Hematol 3, 375–389 (2012).

71. Yan, L., He, Z., Li, W., Liu, N. & Gao, S. The Overexpression of Acyl-CoA Medium-Chain Synthetase-3 (ACSM3) Suppresses the Ovarian Cancer Progression via the Inhibition of Integrin 1/AKT Signaling Pathway. Frontiers in Oncology 11, (2021).

72. Di Narzo, A. F. et al. High-Throughput Identification of the Plasma Proteomic Signature of Inflammatory Bowel Disease. Journal of Crohn’s and Colitis 13, 462–471 (2019).

73. Jacobs, I. et al. Role of Eosinophils in Intestinal Inflammation and Fibrosis in Inflammatory Bowel Disease: An Overlooked Villain? Frontiers in Immunology 12, (2021).

74. Martin, P. et al. Phase I study of the anti-CD74 monoclonal antibody milatuzumab (hLL1) in patients with previously treated B-cell lymphomas. Leuk Lymphoma 56, 3065–3070 (2015).

75. Yao, J. et al. Development and Validation of a Prognostic Gene Signature Correlated With M2 Macrophage Infiltration in Esophageal Squamous Cell Carcinoma. Front Oncol 11, 769727 (2021).

76. Hay, D. et al. Genetic dissection of the a-globin super-enhancer in vivo. Nat Genet 48, 895–903 (2016).

77. Huang, J. et al. Dynamic Control of Enhancer Repertoires Drives Lineage and Stage-Specific Transcription during Hematopoiesis. Dev Cell 36, 9–23 (2016).

78. Bae, S., Park, J. & Kim, J.-S. Cas-OFFinder: a fast and versatile algorithm that searches for potential off-target sites of Cas9 RNA-guided endonucleases. Bioinformatics 30, 1473–1475 (2014).

79. Castro-Mondragon, J. A. et al. JASPAR 2022: the 9th release of the open-access database of transcription factor binding profiles. Nucleic Acids Research 50, D165–D173 (2022).

80. Yukawa, M. et al. AP-1 activity induced by co-stimulation is required for chromatin opening during T cell activation. J Exp Med 217, e20182009 (2020).

81. Perry, M. W., Boettiger, A. N. & Levine, M. Multiple enhancers ensure precision of gap gene-expression patterns in the Drosophila embryo. Proc Natl Acad Sci U S A 108, 13570–13575 (2011).

82. Shin, H. Y. et al. Hierarchy within the mammary STAT5-driven Wap super-enhancer. Nat Genet 48, 904–911 (2016).

83. Huang, J. et al. Dissecting super-enhancer hierarchy based on chromatin interactions. Nat Commun 9, 943 (2018).

84. Snyder, K. J. et al. Inhibition of Bromodomain and Extra Terminal (BET) Domain Activity Modulates the IL-23R/IL-17 Axis and Suppresses Acute Graft-Versus-Host Disease. Front Oncol 11, 760789 (2021).

85. Grubert, F. et al. Landscape of cohesin-mediated chromatin loops in the human genome. Nature 583, 737–743 (2020).

86. Dominguez, A. A. et al. CRISPR-Mediated Synergistic Epigenetic and Transcriptional Control. The CRISPR Journal 5, 264–275 (2022).

87. Schep, A. & University, S. motifmatchr: Fast Motif Matching in R. (2022) doi:10.18129/B9.bioc.motifmatchr.

88. Afgan, E. et al. The Galaxy platform for accessible, reproducible and collaborative biomedical analyses: 2018 update. Nucleic Acids Research 46, W537–W544 (2018).

